# Age and sex differences in lateral ventricle choroid plexus volume in a large, manually segmented, multi-site MRI dataset

**DOI:** 10.64898/2026.11.15.766840

**Authors:** Victor Zeng, Charles J. Wolock, Santiago Estrada, Ling-Yu Huang, Nicolas Raymond, Mohammad Jammoul, Yuan Cao, Matcheri S. Keshavan, Peter J. Fried, David Kuegler, Martin Reuter, Paulo Lizano

**Author notes:** **Corresponding Author**: Paulo Lizano MD, PhD, 300 Crittenden Blvd Rochester NY 14642.

## Abstract

**Background:** The choroid plexus, which produces cerebrospinal fluid and forms the blood–cerebrospinal fluid barrier, is of growing interest in psychiatric and neurodegenerative neuroimaging. However, most volumetric studies have relied on automated segmentation tools that agree poorly with manual tracing, and manually segmented samples have been small, single-site, and narrow in age range.

**Methods:** We analyzed DataPlex, a dataset of 700 manually segmented lateral ventricle choroid plexus labels from 3T T1-weighted MRI scans (ages 6–90 years) drawn from 29 studies and three scanner manufacturers (35 study-by-manufacturer sites). Generalized least squares models accounting for within-site dependence tested associations of age, sex, scanner manufacturer, image resolution, a registration-based image-quality metric, and intracranial volume with choroid plexus and lateral ventricle volumes, including nonlinear (spline) and sex-interaction terms. Exploratory models tested associations between these volumes and cognitive performance.

**Results:** In 699 scans, choroid plexus volume increased linearly with age (63–69 mm³ per decade) and was 56–65 mm³ larger in males than females after covariate adjustment; the age association did not differ by sex. Lateral ventricle volume increased nonlinearly with age, with steeper age-related enlargement in males than females. Scanner manufacturer and image resolution were associated with left lateral ventricle volume; scanner manufacturer was associated with choroid plexus volume only in a sensitivity model. The image-quality metric was not associated with any outcome. In exploratory analyses, larger choroid plexus volume was associated with lower inhibitory control, working memory, and episodic memory scores and with higher emotion recognition scores (uncorrected p < 0.05).

**Conclusions:** In the largest and most diverse manually segmented choroid plexus dataset to date, choroid plexus volume differed by age and sex and was less sensitive than lateral ventricle volume to acquisition differences. These estimates provide reference benchmarks for automated segmentation tools and support adjustment for demographic and technical covariates.

## Introduction

The choroid plexus (ChP) consists of vascularized epithelial tissue lining the lateral, third, and fourth ventricles of the brain, produces cerebrospinal fluid, forms the blood–CSF barrier, and has emerged over the past decade as a structure of substantial interest across psychiatric and neurodegenerative neuroimaging [1–6]. Enlargement of the lateral ventricle ChP (hereafter “ChP”) has been reported across the psychosis spectrum [1], in first-episode antipsychotic-naïve schizophrenia [7], and in individuals at clinical high risk for psychosis [3], where it associates with peripheral inflammatory markers, cognitive impairment, and co-occurring structural brain changes including reduced gray matter and enlarged lateral ventricles (LV). As interest in ChP morphology has grown, so has the recognition that accurately quantifying this small, morphologically complex structure is technically demanding and sensitive to methodological factors, which are often under-examined or unreported in individual studies [8–10].

Age and sex are among the most frequently examined factors that associate with ChP volume, though they are not always reported [11–13]. Cross-sectional studies spanning the adult lifespan find that ChP volume increases, linearly with age in one deep-learning-validated cohort [11], and nonlinearly across a dataset with ages 21–94 [12], and also a 641-participant HCP-Aging dataset [13]. Sex effects are considerably less consistent with some cohorts reporting significantly larger ChP volume in males [11], while others find no significant cross-sectional sex difference despite sex differences in ChP perfusion and diffusivity metrics [12]. At least one manually segmented clinical sample found an apparent larger ChP volume in males that disappeared after adjustment for age and intracranial volume [7]. Longitudinally, in a study of cognitive decline, women showed roughly double the annual rate of ChP growth compared with men [14], suggesting age and sex may interact rather than act independently.

A key methodological issue across these studies is how ChP volume is measured. Automated pipelines, chiefly FreeSurfer, are the only approaches that have scaled to large or lifespan-spanning samples (e.g., n=613–5,551) [14,15], but direct comparisons against manual tracing show these automated labels are unreliable for the ChP. FreeSurfer segmentations agreed poorly with manual ground truth across a multi-vendor, 8-site sample (Dice ≈0.29) [3]. Even within a single site and scanner, FreeSurfer-derived volumes correlated only moderately with manual volumes (r=0.659) [16]. Deep-learning approaches narrow this gap (Dice 0.72–0.74 against manual ground truth [11]), but their accuracy remains limited, and they have been validated mostly in older adults. Manual tracing or manual corrections of FreeSurfer predictions has therefore served as the reference standard used in every study that has directly compared methods, yet manually segmented ChP cohorts to date are uniformly small (<200 scans) and restricted to narrow, adult-only age ranges or infants [17]. This tradeoff compounds with other under-examined sources of bias. Scanner field strength, vendor, and image resolution can also bias structural and ventricular volume estimates [18,19]. Unaddressed motion artifact and image-quality variability can further bias automated volumetric estimates and can generate both false-positive and false-negative structure–behavior associations in large samples [20–22]. Manual segmentation with rigorous quality control remains the most defensible approach for a structure this small and this consequential, but no prior study has applied it at scale across a demographically and technically diverse sample.

ChP volume is also structurally, and plausibly mechanistically, intertwined with LV volume. LV enlargement is the structural MRI correlate most consistently associated with ChP enlargement (r=0.38) [7], with the two structures showing increased joint variance in schizophrenia-spectrum disorders [16]. Thus, larger ventricles could plausibly bias ChP volume estimates given their shared boundary at the choroidal fissure, though automated segmentation performance has been reported as stable across a range of LV volumes [11], a claim not yet established for manual tracing. No published dataset, to our knowledge, has manually segmented ChP volume in a sample both large and demographically and technically heterogeneous enough to disentangle these effects. Here, we manually segmented the lateral ventricle ChP in 700 brain MRI scans spanning a wide age range from childhood through late life, biological sex, and multiple scanner types, enabling us to characterize the independent contributions of these demographic and technical factors to ChP volume and to compare them with their effects on LV volume. Given this literature, we hypothesized that ChP volume would show significant, independent associations with both age and sex across our sample, and that these demographic effects would remain detectable once scanner manufacturer, image resolution, image quality, and intracranial volume were explicitly accounted for. Testing these hypotheses in a manually segmented sample spanning this range of age, sex, and scanner types allows us to determine whether the age and sex differences reported in smaller, more homogeneous, and largely automated-segmentation cohorts generalize across the full developmental and aging spectrum, and to establish reference benchmarks for the demographic and technical covariates that future ChP volumetric research should account for.

## Materials and methods

### Data overview

DataPlex comprises manually segmented lateral ventricle ChP volumes from 700 3-Tesla T1-weighted MRI scans. Manual segmentations were conducted by trained raters (mean Dice coefficient (DC) across raters = 0.81; minimum = 0.75; maximum = 0.86) and verified through a formal quality-control process by expert raters (mean DC = 0.89; minimum = 0.84; maximum = 0.92). T2-weighted images acquired within-session were available for half (N=351) of the DataPlex sample, and the T2-highlighted vasculature provided external validation for our segmentation quality (mean DC with vs without T2 = 0.95; maximum = 0.98; minimum = 0.84). Scans were selected to vary along three dimensions: (1) MRI scanner manufacturer, (2) image resolution (mm), and (3) demographic characteristics, including binned age group, sex, and phenotype. Further details on DataPlex’s selection criteria are provided in Huang et al. (2026) [23]. The categories for MRI scanner manufacturers included General Electric (GE), Philips, and Siemens, and there were three isotropic image resolutions the MRI scans were acquired in: 0.7 mm, 0.8 mm, and 1.0 mm.

DataPlex was constructed by stratified sampling of 29 open-source and investigator-contributed studies. These contributing studies obtained consent from participants for their data to be shared publicly. The University of Rochester Medical Center Institutional Review Board (IRB) reviewed the present secondary analysis of de-identified data and waived the requirement for additional informed consent; all contributing studies obtained written informed consent from adult participants, and written parental permission with child assent for minors, under their local IRBs, and we established Data Use Agreements (DUA) with the contributing data repositories and sites since MRI is categorized as sensitive data. Sources included the NIMH Data Archives, OpenNeuro, and Beth Israel Deaconess Medical Center. Acquisition parameters and data use agreements are described in S1 File.

The studies contributing to the 700 DataPlex scans varied in MRI scanner and acquisition protocol both between and within studies. Herein we define “site” as the combined identifier of the research study and the MRI scanner manufacturer. For example, the Adolescent Brain Cognitive Development project (ABCD) used three scanner manufacturers: Siemens, GE, and Philips. Rather than treating ABCD as a single site or assigning each of its contributing research sites in its own category, we identified scans from ABCD as “ABCD-Siemens,” “ABCD-GE,” and “ABCD-Philips.” Within a given site, scans may still differ in scanner model (e.g., GE Signa vs. GE MR750), scanner software version, head coil channel count, or acquisition location; because this information was not consistently available across contributing studies, these finer differences were not modeled in the analysis.

Beyond MRI, several contributing sites administered a range of cognitive assessments, which were used to explore relationships with ChP and LV volumetrics. These include the PENN-40 [24], a test of emotional recognition, and the NIH Toolbox cognitive battery, comprising the Flanker Inhibitory Control and Attention Test, Dimensional Change Card Sort Test, List Sorting Working Memory Test, Oral Reading Recognition Test, Picture Sequence Memory Test, and Picture Vocabulary Test. **Table 1** lists all examinations used in this analysis along with the specific measure drawn from each.

**Table 1.**
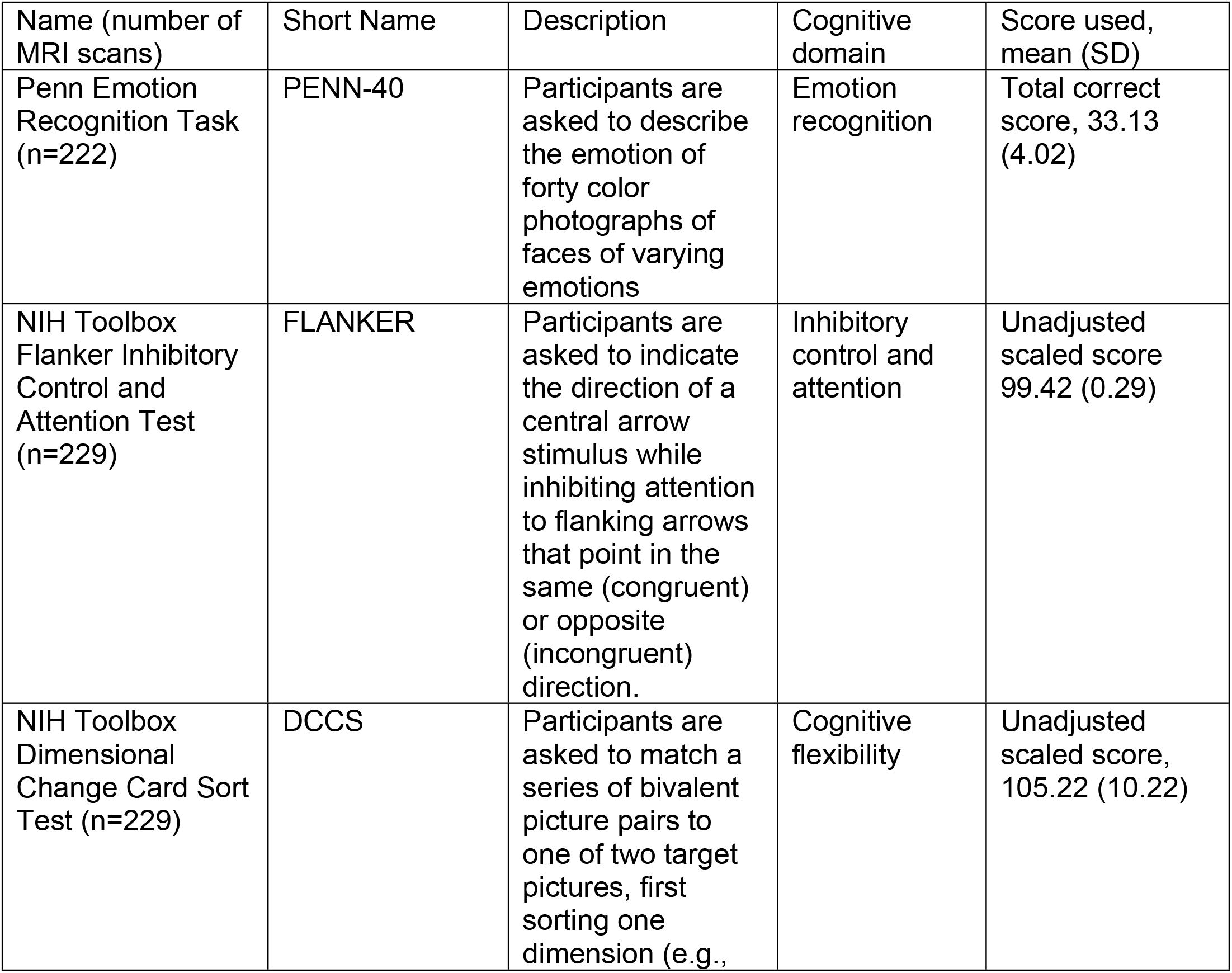
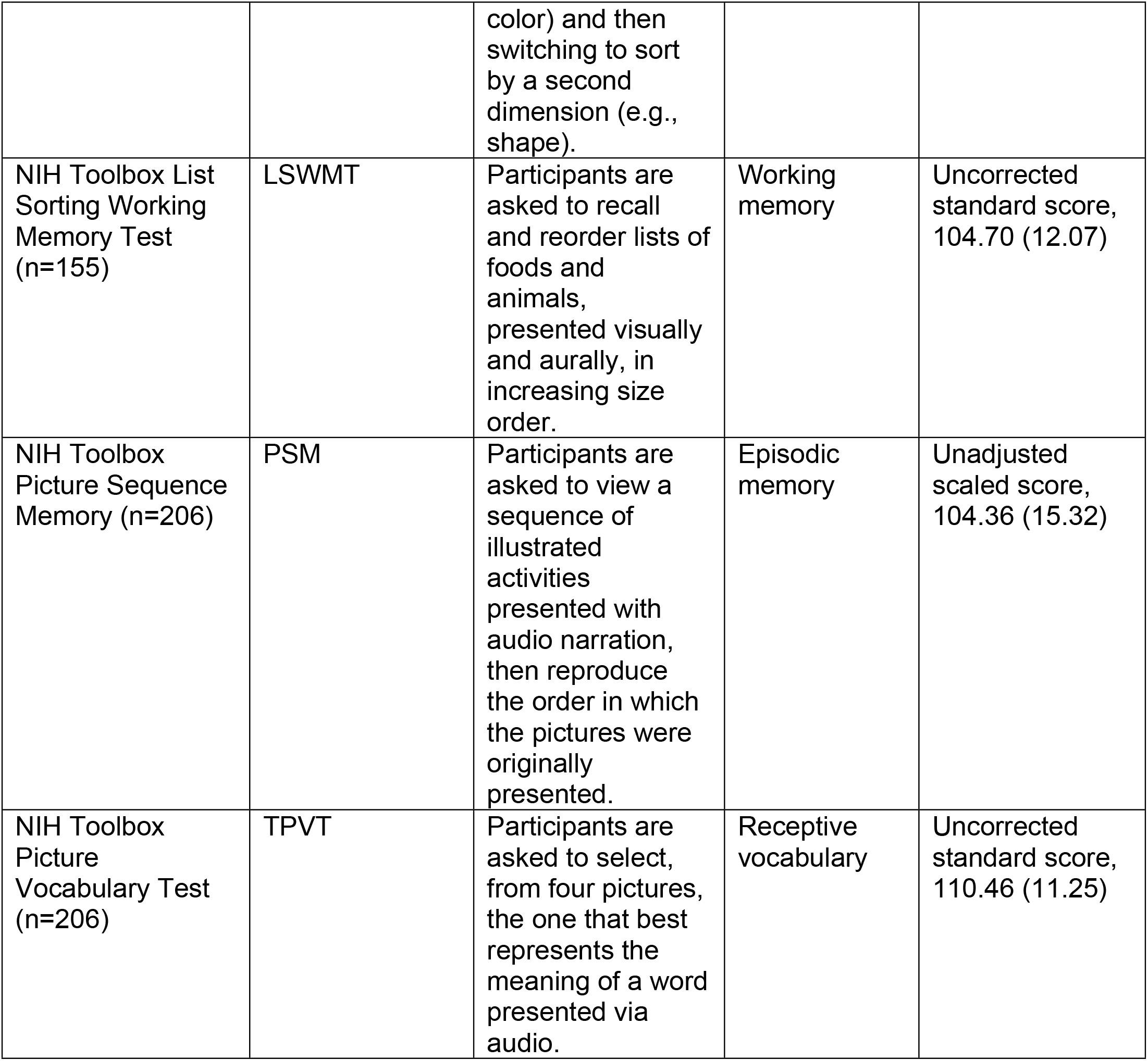
Cognitive measures used in the analysis.

### Data processing

Manual ChP segmentations were conducted using 3D Slicer (version 5.2.2, www.slicer.org) [25] and saved in segmentation labelmap (.seg.nrrd) format. 3D Slicer’s built-in data converter was then used to convert the .seg.nrrd label maps to the .mgz format readable by FreeSurfer. FreeSurfer 7.1.0 [26] was then used to extract volumetric measurements of the manually segmented left and right ChP from these files. Throughout this study, “left and right ChP” refers specifically to the ChP volume located within the LV.

To assess the image quality of the structural scans, the FSQC tool was used to calculate a range of MRI quality-control (QC) metrics [21,27]. Two QC metrics were of particular interest in this analysis as potential predictors of image quality: “rot_tal_x” and “gm_snr_orig.” The rot_tal_x quantifies the degree of rotation applied during Talairach registration, and it can be used to flag failed registrations due to possible non-standard head positioning or anatomy. The gm_snr_orig summarizes the gray matter signal-to-noise ratio which quantifies the scan’s ability to resolve tissue layers. To examine the relationships between ChP volume and ventricular/whole-brain volumes, all 700 T1-weighted scans were processed using FastSurfer v2.5.4 [28,29] to extract measurements for estimated total intracranial volume (eTIV) and left and right LV volumes. One scan from the DataPlex sample failed FastSurfer’s surface reconstruction module, and it was excluded from QC metric extraction and subsequent analysis.

FastSurfer also extracted and reported estimates for left and right lateral ventricle ChP, but it occasionally mislabeled ChP tissue as LV due to incorrect boundary definition. This created inconsistency in LV volume estimates to be analyzed against manual segmented ChP volumes. To address this, we performed a manual correction procedure by summing FastSurfer-derived LV and ChP volume estimates (which recovered the full volume enclosed by the LV outer boundaries) and then subtracting the manually segmented ChP volume. This calculation was performed separately for the left and right LV to provide a better estimate of LV volume.

### Statistical analysis

Statistical analyses were conducted in R 4.4.2 using RStudio. To account for potential site-specific effects on ChP volume, generalized least squares (GLS) models were fitted using the Gls function from the rms package; every GLS model incorporated a compound symmetry correlation structure (corCompSymm), implemented via the *nlme* package, to account for the non-independence of observations within site categories.

To examine the associations of several predictors of interest with ChP and LV volumes, ChP and LV volume estimates were each included as the outcome in a separate GLS model. Predictors included were age, sex, scanner manufacturer, image resolution, rot_tal_x, and eTIV. Age, eTIV, and rot_tal_x were modeled with restricted cubic splines (three knots) to test potential non-linear associations with each outcome. For predictors represented by more than one coefficient (spline terms and three-level categorical predictors), p values are from joint Wald tests of all coefficients for that predictor; in interaction models, the p value for a main effect also includes its interaction terms. Pair-wise post hoc comparisons are corrected using the false discovery rate (q). Statistical significance was set at two-sided α = 0.05. A parallel model using linear terms with the same predictors was also fit to report linear associations between the predictors and ChP or LV volumes. A sensitivity model substituting gm_snr_orig for rot_tal_x is reported in S1 File (Table B and Figs A–D).

To examine potential relationships between the cognitive examinations and ChP and LV volume, each cognitive measure was modeled as the outcome in a linear (non-spline) GLS model. Predictors for this analysis included ChP or LV volume, age, sex, scanner manufacturer, image resolution, rot_tal_x, and eTIV. These cognitive analyses were exploratory; p values were not corrected for multiple comparisons and should be interpreted accordingly. Image resolution was omitted from the PENN-40 models to avoid collinearity, because a single study contributed all Philips and all 1.0 mm PENN-40 scans (S1 File).

## Results

The final analytic sample consisted of 699 manually segmented MRI scans (one scan was excluded for failing FastSurfer’s surface reconstruction module), spanning 35 sites (Table A in S1 File). Participants ranged broadly in age (mean 39.1 ± 23.7 years) and were nearly evenly split by sex (342 male, 357 female). Scans were acquired across three scanner manufacturers, GE (n=251), Philips (n=196), and Siemens (n=252), and three image resolutions: 0.7 mm (n=22), 0.8 mm (n=329), and 1.0 mm (n=348). Mean estimated total intracranial volume (eTIV) was 1,513,927 ± 187,771 mm³, mean gray matter signal-to-noise ratio was 5.46 ± 1.69, and mean Talairach x-axis rotation was −0.21 ± 0.12. Mean ChP volume was 867.6 ± 365.6 mm³ on the left and 849.2 ± 347.0 mm³ on the right; mean LV volume was 9,184.9 ± 6,505.1 mm³ on the left and 8,284.5 ± 5,704.5 mm³ on the right. Full demographic and technical characteristics are reported in **Table 2**.

**Table 2.** Demographic and technical characteristics of the analytic sample (n=699).

|  |  |
|---|---|
| Age (range, mean, SD) | 6.3 – 89.8 \| 39.1 (23.7) |
| Sex (M, F) | 342 \| 357 |
| Gray Matter Signal to Noise Ratio (mean, SD) | 5.46 (1.69) |
| Rotation of the Talairach in the x Axis (mean, SD) | -0.21 (0.12) |
| Image Resolution (0.7, 0.8, 1.0 mm isotropic) | 22 \| 329 \| 348 |
| Scanner Manufacturer (GE, Philips, Siemens) | 251 \| 196 \| 252 |
| Estimated Total Intracranial Volume, mm<sup>3</sup> (mean, SD) | 1,513,927 (187,771) |
| Left ChP, mm<sup>3</sup> (mean, SD) | 867.6 (365.6) |
| Right ChP, mm<sup>3</sup> (mean, SD) | 849.2 (347.0) |
| Left LV, mm<sup>3</sup> (mean, SD) | 9184.9 (6505.1) |
| Right LV, mm<sup>3</sup> (mean, SD) | 8284.5 (5704.5) |

**Notes:** ChP, choroid plexus; LV, lateral ventricle; SD, standard deviation; M, male; F, female; mm, millimeter; GE, General Electric

### Results of the GLS models for ChP and LV volume

**Table 3** summarizes GLS outcome statistics from restricted cubic splines using linear and non-linear terms of age, x-axis rotation angle, and eTIV as predictors. The estimated β coefficients, confidence intervals, and p-values are reported for the overall sample.

**Table 3.**
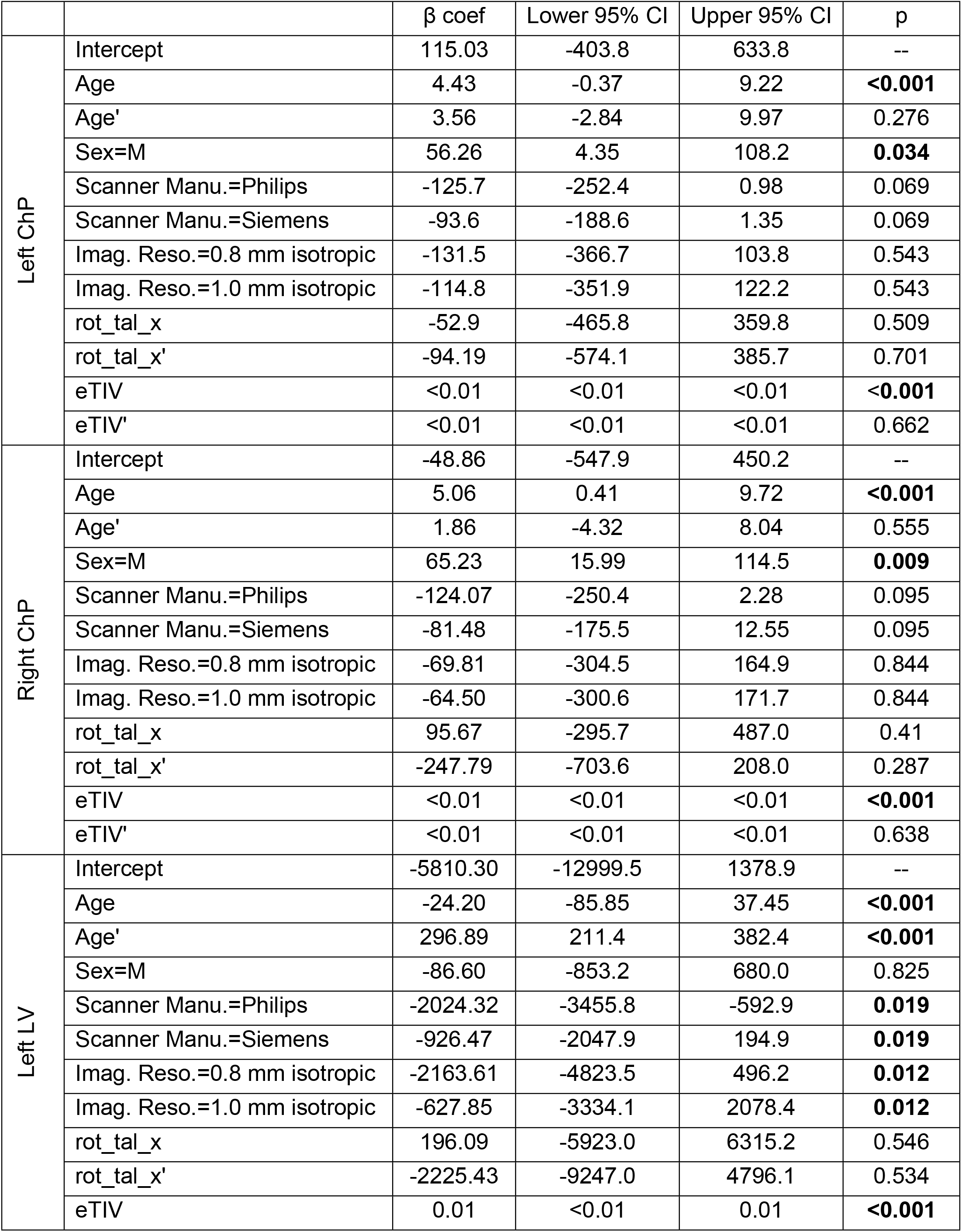
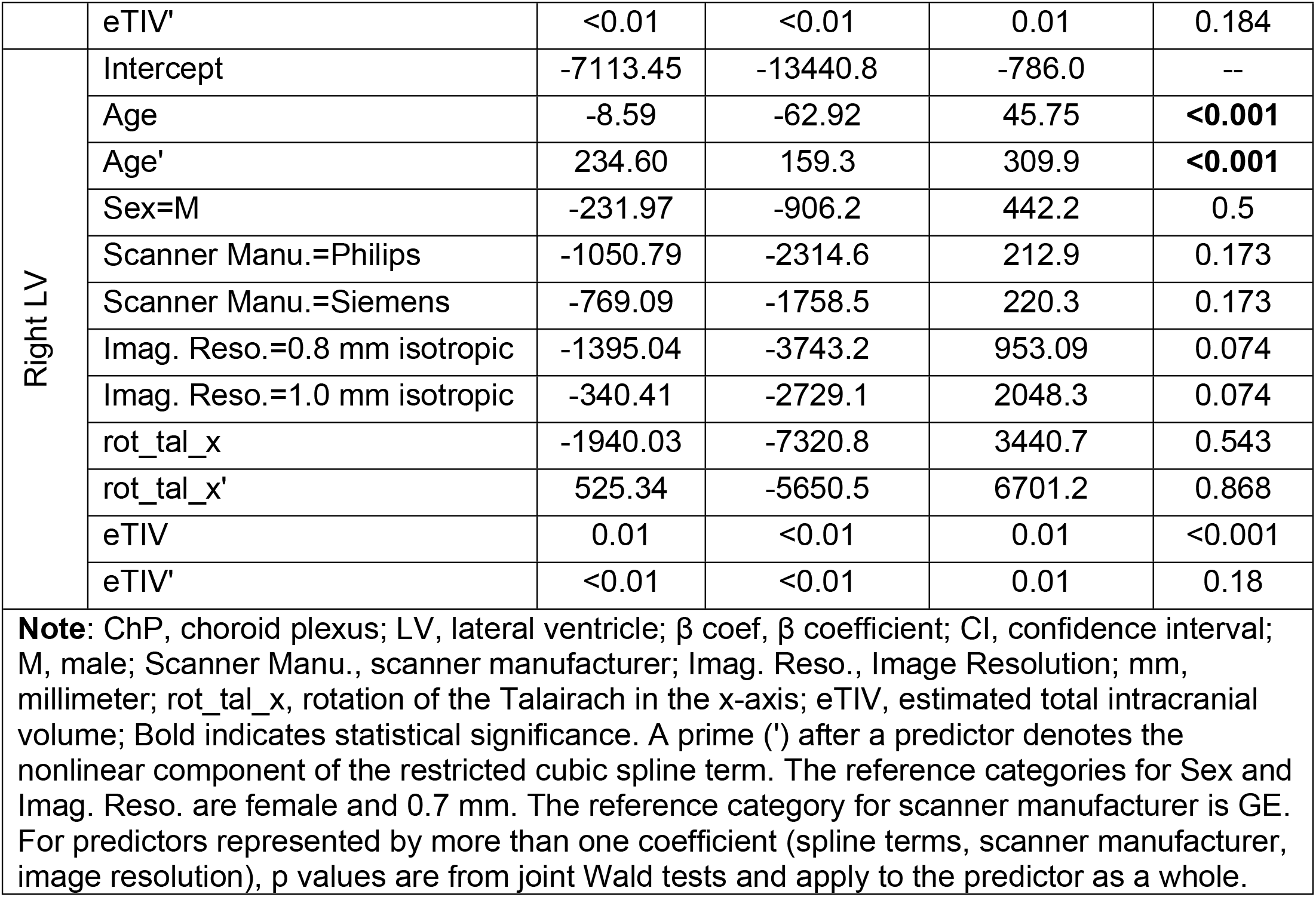
GLS models with restricted cubic splines for left and right ChP and LV volumes.

|  |  | β coef | Lower 95% CI | Upper 95% CI | p |
|---|---|---|---|---|---|
| Left ChP | Intercept | 115.03 | -403.8 | 633.8 | -- |
|  | Age | 4.43 | -0.37 | 9.22 | **<0.001** |
|  | Age' | 3.56 | -2.84 | 9.97 | 0.276 |
|  | Sex=M | 56.26 | 4.35 | 108.2 | **0.034** |
|  | Scanner Manu.=Philips | -125.7 | -252.4 | 0.98 | 0.069 |
|  | Scanner Manu.=Siemens | -93.6 | -188.6 | 1.35 | 0.069 |
|  | Imag. Reso.=0.8 mm isotropic | -131.5 | -366.7 | 103.8 | 0.543 |
|  | Imag. Reso.=1.0 mm isotropic | -114.8 | -351.9 | 122.2 | 0.543 |
|  | rot_tal_x | -52.9 | -465.8 | 359.8 | 0.509 |
|  | rot_tal_x' | -94.19 | -574.1 | 385.7 | 0.701 |
|  | eTIV | <0.01 | <0.01 | <0.01 | <**0.001** |
|  | eTIV' | <0.01 | <0.01 | <0.01 | 0.662 |
| Right ChP | Intercept | -48.86 | -547.9 | 450.2 | -- |
|  | Age | 5.06 | 0.41 | 9.72 | **<0.001** |
|  | Age' | 1.86 | -4.32 | 8.04 | 0.555 |
|  | Sex=M | 65.23 | 15.99 | 114.5 | **0.009** |
|  | Scanner Manu.=Philips | -124.07 | -250.4 | 2.28 | 0.095 |
|  | Scanner Manu.=Siemens | -81.48 | -175.5 | 12.55 | 0.095 |
|  | Imag. Reso.=0.8 mm isotropic | -69.81 | -304.5 | 164.9 | 0.844 |
|  | Imag. Reso.=1.0 mm isotropic | -64.50 | -300.6 | 171.7 | 0.844 |
|  | rot_tal_x | 95.67 | -295.7 | 487.0 | 0.41 |
|  | rot_tal_x' | -247.79 | -703.6 | 208.0 | 0.287 |
|  | eTIV | <0.01 | <0.01 | <0.01 | **<0.001** |
|  | eTIV' | <0.01 | <0.01 | <0.01 | 0.638 |
| Left LV | Intercept | -5810.30 | -12999.5 | 1378.9 | -- |
|  | Age | -24.20 | -85.85 | 37.45 | **<0.001** |
|  | Age' | 296.89 | 211.4 | 382.4 | **<0.001** |
|  | Sex=M | -86.60 | -853.2 | 680.0 | 0.825 |
|  | Scanner Manu.=Philips | -2024.32 | -3455.8 | -592.9 | **0.019** |
|  | Scanner Manu.=Siemens | -926.47 | -2047.9 | 194.9 | **0.019** |
|  | Imag. Reso.=0.8 mm isotropic | -2163.61 | -4823.5 | 496.2 | **0.012** |
|  | Imag. Reso.=1.0 mm isotropic | -627.85 | -3334.1 | 2078.4 | **0.012** |
|  | rot_tal_x | 196.09 | -5923.0 | 6315.2 | 0.546 |
|  | rot_tal_x' | -2225.43 | -9247.0 | 4796.1 | 0.534 |
|  | eTIV | 0.01 | <0.01 | 0.01 | **<0.001** |
|  | eTIV' | <0.01 | <0.01 | 0.01 | 0.184 |
| Right LV | Intercept | -7113.45 | -13440.8 | -786.0 | -- |
|  | Age | -8.59 | -62.92 | 45.75 | **<0.001** |
|  | Age' | 234.60 | 159.3 | 309.9 | **<0.001** |
|  | Sex=M | -231.97 | -906.2 | 442.2 | 0.5 |
|  | Scanner Manu.=Philips | -1050.79 | -2314.6 | 212.9 | 0.173 |
|  | Scanner Manu.=Siemens | -769.09 | -1758.5 | 220.3 | 0.173 |
|  | Imag. Reso.=0.8 mm isotropic | -1395.04 | -3743.2 | 953.09 | 0.074 |
|  | Imag. Reso.=1.0 mm isotropic | -340.41 | -2729.1 | 2048.3 | 0.074 |
|  | rot_tal_x | -1940.03 | -7320.8 | 3440.7 | 0.543 |
|  | rot_tal_x' | 525.34 | -5650.5 | 6701.2 | 0.868 |
|  | eTIV | 0.01 | <0.01 | 0.01 | <0.001 |
|  | eTIV' | <0.01 | <0.01 | 0.01 | 0.18 |

**Note**: ChP, choroid plexus; LV, lateral ventricle; β coef, β coefficient; CI, confidence interval; M, male; Scanner Manu., scanner manufacturer; Imag. Reso., Image Resolution; mm, millimeter; rot\_tal\_x, rotation of the Talairach in the x-axis; eTIV, estimated total intracranial volume; Bold indicates statistical significance. A prime (') after a predictor denotes the nonlinear component of the restricted cubic spline term. The reference categories for Sex and Imag. Reso. are female and 0.7 mm. The reference category for scanner manufacturer is GE. For predictors represented by more than one coefficient (spline terms, scanner manufacturer, image resolution), p values are from joint Wald tests and apply to the predictor as a whole.

**Table 4** reports the same predictors and outcomes as **Table 3**, without the spline non-linear terms in the model.

**Table 4.** Linear GLS models for left and right ChP and LV volumes.

|  |  | β coef | Lower 95% CI | Upper 95% CI | p |
|---|---|---|---|---|---|
| Left ChP | Intercept | -54.12 | -379.2 | 270.9 | -- |
|  | Age | 6.93 | 5.32 | 8.53 | **<0.001** |
|  | Sex=M | 56.82 | 5.02 | 108.6 | **0.032** |
|  | Scanner Manu.=Philips | -122.2 | -248.9 | 4.50 | 0.079 |
|  | Scanner Manu.=Siemens | -91.21 | -186.0 | 3.61 | 0.079 |
|  | Imag. Reso.=0.8 mm isotropic | -118.2 | -353.0 | 116.6 | 0.601 |
|  | Imag. Reso.=1.0 mm isotropic | -99.59 | -335.7 | 136.5 | 0.601 |
|  | rot_tal_x | -115.3 | -332.3 | 101.7 | 0.298 |
|  | eTIV | <0.01 | <0.01 | <0.01 | **<0.001** |
| Right ChP | Intercept | -62.40 | -377.5 | 252.7 | -- |
|  | Age | 6.30 | 4.74 | 7.85 | **<0.001** |
|  | Sex=M | 64.27 | 15.13 | 113.4 | **0.01** |
|  | Scanner Manu.=Philips | -117.7 | -242.4 | 6.99 | 0.115 |
|  | Scanner Manu.=Siemens | -76.47 | -169.3 | 16.35 | 0.115 |
|  | Imag. Reso.=0.8 mm isotropic | -65.67 | -296.9 | 165.5 | 0.855 |
|  | Imag. Reso.=1.0 mm isotropic | -58.12 | -290.4 | 174.1 | 0.855 |
|  | rot_tal_x | -87.11 | -293.4 | 119.1 | 0.408 |
|  | eTIV | <0.01 | <0.01 | <0.01 | **<0.001** |
| Left LV | Intercept | -15809.5 | -20837.5 | -10781.5 | -- |
|  | Age | 188.6 | 163.8 | 213.5 | **<0.001** |
|  | Sex=M | -92.42 | -882.7 | 697.9 | 0.819 |
|  | Scanner Manu.=Philips | -1815.5 | -3794.4 | 163.5 | 0.194 |
|  | Scanner Manu.=Siemens | -699.3 | -2175.1 | 776.5 | 0.194 |
|  | Imag. Reso.=0.8 mm isotropic | -974.6 | -4643.2 | 2694.0 | 0.081 |
|  | Imag. Reso.=1.0 mm isotropic | 694.2 | -2992.2 | 4380.6 | 0.081 |
|  | rot_tal_x | -737.9 | -4052.6 | 2576.6 | 0.663 |
|  | eTIV | 0.01 | 0.01 | 0.01 | **<0.001** |
| Right LV | Intercept | -14816.6 | -19160.6 | -10472.6 | -- |
|  | Age | 158.8 | 137.4 | 180.3 | **<0.001** |
|  | Sex=M | -233.7 | -925.1 | 457.7 | 0.508 |
|  | Scanner Manu.=Philips | -923.8 | -2617.9 | 770.4 | 0.48 |
|  | Scanner Manu.=Siemens | -615.7 | -1883.2 | 651.9 | 0.48 |
|  | Imag. Reso.=0.8 mm isotropic | -412.7 | -3552.9 | 2727.5 | 0.187 |
|  | Imag. Reso.=1.0 mm isotropic | 759.1 | -2398.2 | 3916.3 | 0.187 |
|  | rot_tal_x | -987.1 | -3883.9 | 1909.7 | 0.504 |
|  | eTIV | 0.01 | 0.01 | 0.01 | **<0.001** |

**Note**: ChP, choroid plexus; LV, lateral ventricle; β coef, β coefficient; CI, confidence interval; M, male; Scanner Manu., scanner manufacturer; Imag. Reso., Image Resolution; mm, millimeter; rot_tal_x, rotation of the talairach in the x-axis; eTIV, estimated total intracranial volume; Bold indicates statistical significance. The reference categories for Sex and Imag. Reso. are female and 0.7 mm. The reference category for scanner manufacturer is GE. For predictors represented by more than one coefficient (spline terms, scanner manufacturer, image resolution), p values are from joint Wald tests and apply to the predictor as a whole.

Finally, **Table 5** reports the same predictors and outcomes, in a linear model, but reported sex interactions with age and eTIV in the model

**Table 5.**
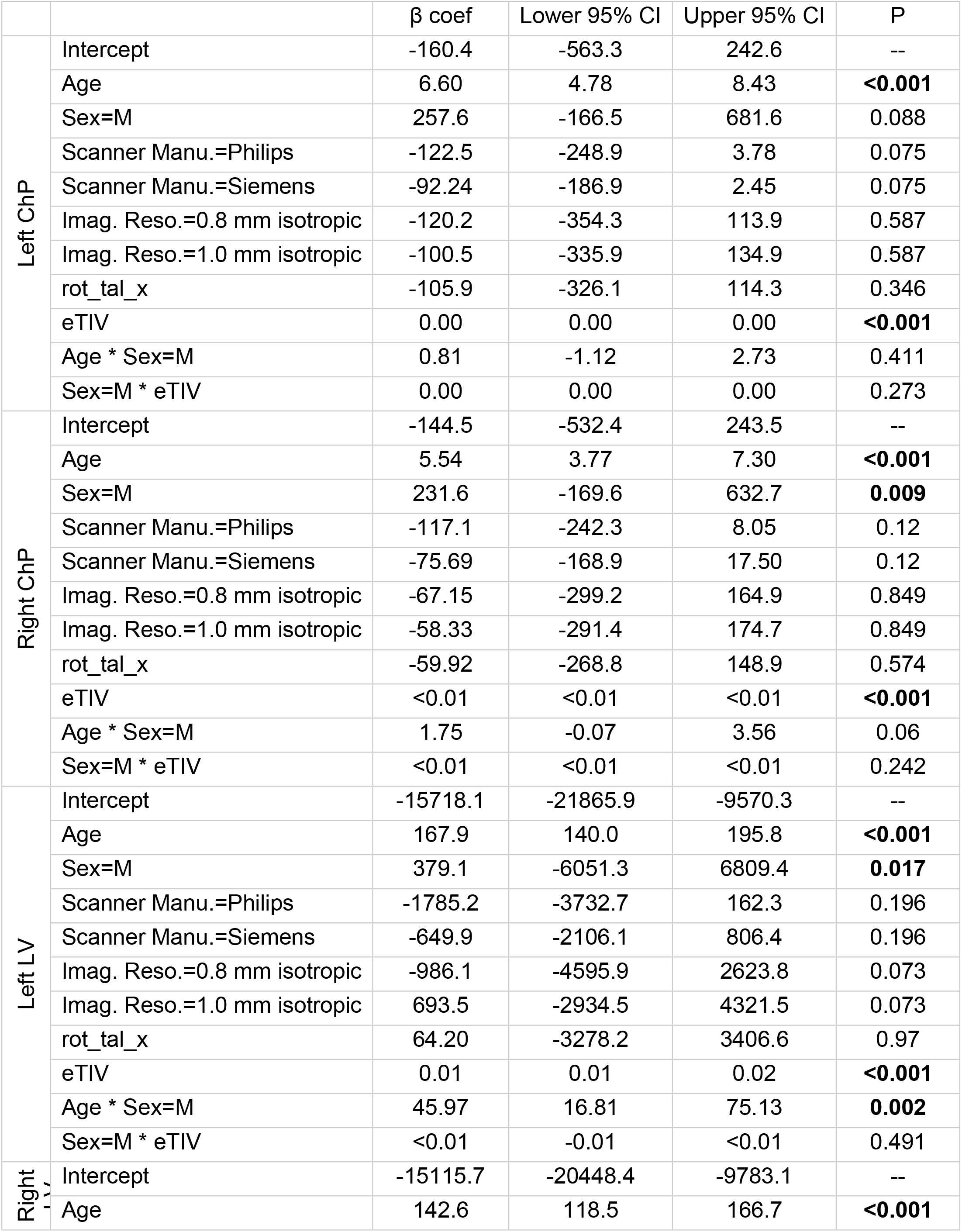
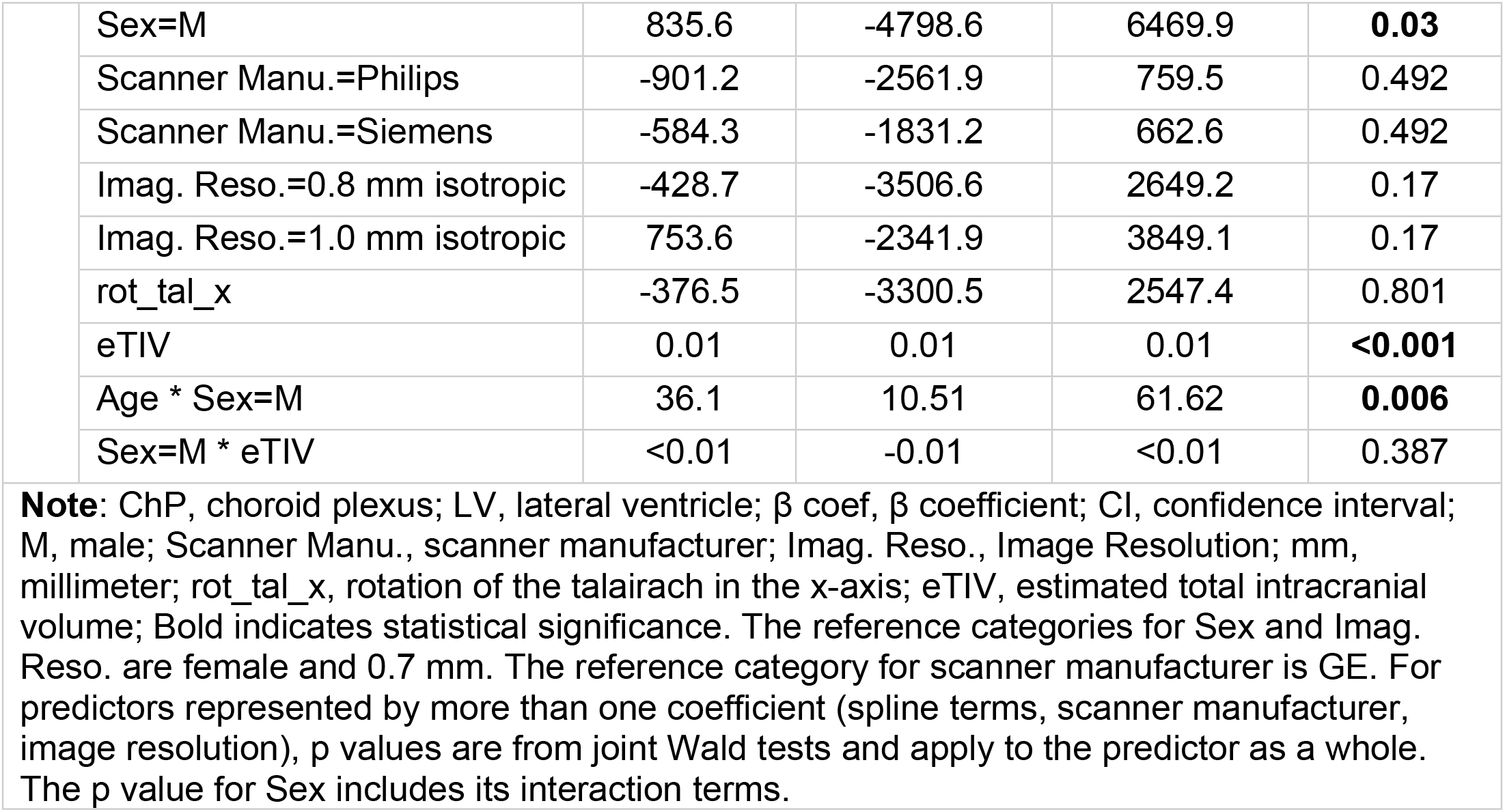
Linear GLS models with sex interaction terms for left and right ChP and LV volumes.

|  |  | β coef | Lower 95% CI | Upper 95% CI | P |
|---|---|---|---|---|---|
| Left ChP | Intercept | -160.4 | -563.3 | 242.6 | -- |
|  | Age | 6.60 | 4.78 | 8.43 | **<0.001** |
|  | Sex=M | 257.6 | -166.5 | 681.6 | 0.088 |
|  | Scanner Manu.=Philips | -122.5 | -248.9 | 3.78 | 0.075 |
|  | Scanner Manu.=Siemens | -92.24 | -186.9 | 2.45 | 0.075 |
|  | Imag. Reso.=0.8 mm isotropic | -120.2 | -354.3 | 113.9 | 0.587 |
|  | Imag. Reso.=1.0 mm isotropic | -100.5 | -335.9 | 134.9 | 0.587 |
|  | rot_tal_x | -105.9 | -326.1 | 114.3 | 0.346 |
|  | eTIV | 0.00 | 0.00 | 0.00 | **<0.001** |
|  | Age * Sex=M | 0.81 | -1.12 | 2.73 | 0.411 |
|  | Sex=M * eTIV | 0.00 | 0.00 | 0.00 | 0.273 |
| Right ChP | Intercept | -144.5 | -532.4 | 243.5 | -- |
|  | Age | 5.54 | 3.77 | 7.30 | **<0.001** |
|  | Sex=M | 231.6 | -169.6 | 632.7 | **0.009** |
|  | Scanner Manu.=Philips | -117.1 | -242.3 | 8.05 | 0.12 |
|  | Scanner Manu.=Siemens | -75.69 | -168.9 | 17.50 | 0.12 |
|  | Imag. Reso.=0.8 mm isotropic | -67.15 | -299.2 | 164.9 | 0.849 |
|  | Imag. Reso.=1.0 mm isotropic | -58.33 | -291.4 | 174.7 | 0.849 |
|  | rot_tal_x | -59.92 | -268.8 | 148.9 | 0.574 |
|  | eTIV | <0.01 | <0.01 | <0.01 | **<0.001** |
|  | Age * Sex=M | 1.75 | -0.07 | 3.56 | 0.06 |
|  | Sex=M * eTIV | <0.01 | <0.01 | <0.01 | 0.242 |
| Left LV | Intercept | -15718.1 | -21865.9 | -9570.3 | -- |
|  | Age | 167.9 | 140.0 | 195.8 | **<0.001** |
|  | Sex=M | 379.1 | -6051.3 | 6809.4 | **0.017** |
|  | Scanner Manu.=Philips | -1785.2 | -3732.7 | 162.3 | 0.196 |
|  | Scanner Manu.=Siemens | -649.9 | -2106.1 | 806.4 | 0.196 |
|  | Imag. Reso.=0.8 mm isotropic | -986.1 | -4595.9 | 2623.8 | 0.073 |
|  | Imag. Reso.=1.0 mm isotropic | 693.5 | -2934.5 | 4321.5 | 0.073 |
|  | rot_tal_x | 64.20 | -3278.2 | 3406.6 | 0.97 |
|  | eTIV | 0.01 | 0.01 | 0.02 | **<0.001** |
|  | Age * Sex=M | 45.97 | 16.81 | 75.13 | **0.002** |
|  | Sex=M * eTIV | <0.01 | -0.01 | <0.01 | 0.491 |
| Right [illegible] | Intercept | -15115.7 | -20448.4 | -9783.1 | -- |
|  | Age | 142.6 | 118.5 | 166.7 | **<0.001** |
|  | Sex=M | 835.6 | -4798.6 | 6469.9 | **0.03** |
|  | Scanner Manu.=Philips | -901.2 | -2561.9 | 759.5 | 0.492 |
|  | Scanner Manu.=Siemens | -584.3 | -1831.2 | 662.6 | 0.492 |
|  | Imag. Reso.=0.8 mm isotropic | -428.7 | -3506.6 | 2649.2 | 0.17 |
|  | Imag. Reso.=1.0 mm isotropic | 753.6 | -2341.9 | 3849.1 | 0.17 |
|  | rot_tal_x | -376.5 | -3300.5 | 2547.4 | 0.801 |
|  | eTIV | 0.01 | 0.01 | 0.01 | **<0.001** |
|  | Age * Sex=M | 36.1 | 10.51 | 61.62 | **0.006** |
|  | Sex=M * eTIV | <0.01 | -0.01 | <0.01 | 0.387 |

**Note**: ChP, choroid plexus; LV, lateral ventricle; β coef, β coefficient; CI, confidence interval; M, male; Scanner Manu., scanner manufacturer; Imag. Reso., Image Resolution; mm, millimeter; rot_tal_x, rotation of the talairach in the x-axis; eTIV, estimated total intracranial volume; Bold indicates statistical significance. The reference categories for Sex and Imag. Reso. are female and 0.7 mm. The reference category for scanner manufacturer is GE. For predictors represented by more than one coefficient (spline terms, scanner manufacturer, image resolution), p values are from joint Wald tests and apply to the predictor as a whole. The p value for Sex includes its interaction terms.

Left and right ChP volumes were both related to age (p<0.001), with no significant nonlinear component for either (p=0.276 and p=0.555, respectively) when modeled using restricted cubic splines (Figs 1 and 2; Tables 3 and 4). LV volume was also related to age; however, the nonlinear component was significant for both the left and right LV (both p<0.001), indicating a nonlinear relationship between age and LV volume (Figs 3 and 4; Tables 3 and 4). Under a linear model, adjusting for all other covariates, average left ChP volume increases by 69.28 mm³ (95% CI: 53.21–85.34) per 10-year difference in age, and average right ChP volume increases by 62.96 mm³ (95% CI: 47.40–78.52) per 10-year difference in age. In the linear model, the average left LV increases by 1886.15 mm³ (95% CI: 1637.75–2134.55) and the average right LV increases by 1588.30 mm³ (95% CI: 1373.60–1803.00) per 10-year difference in age.

**Fig 1.**
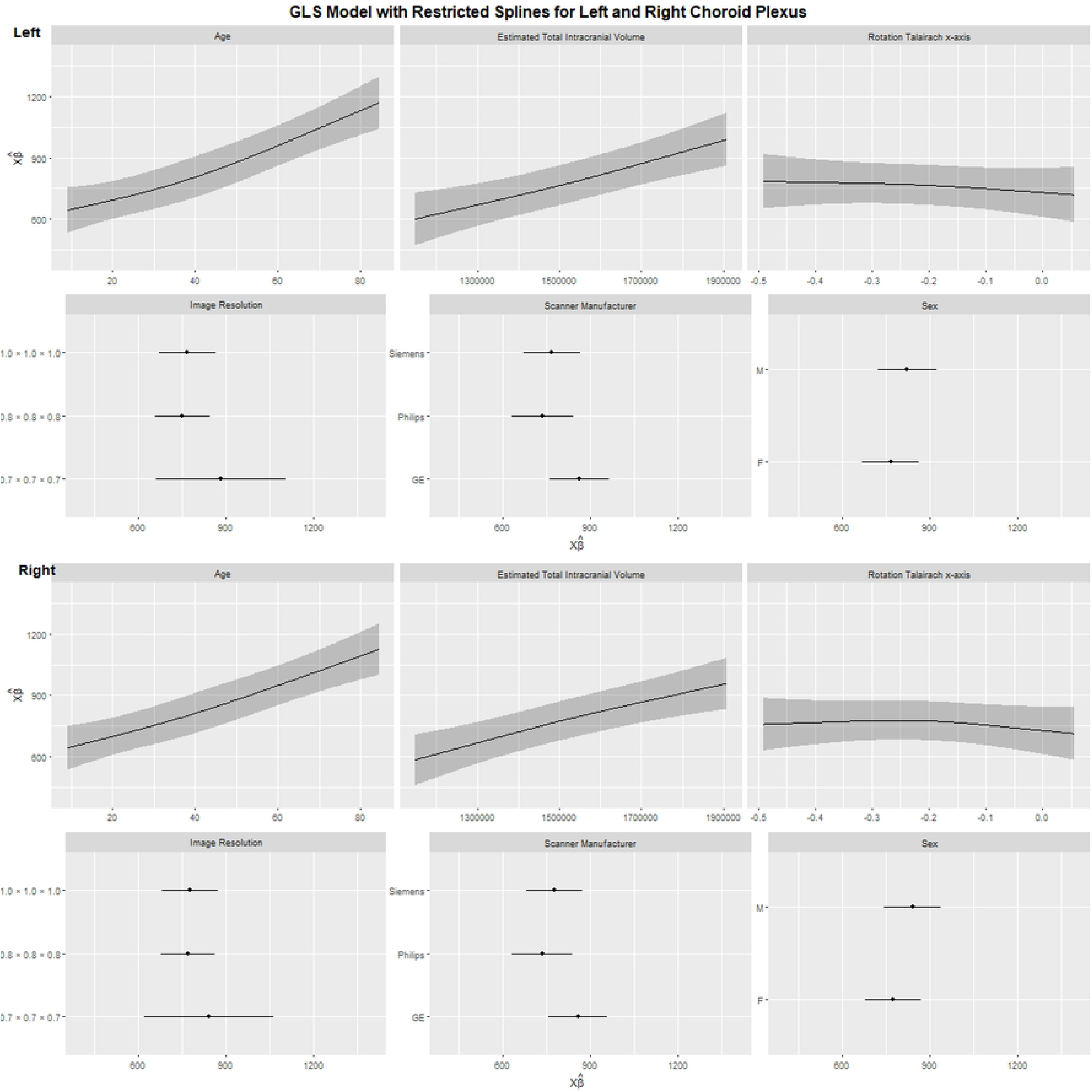
Choroid plexus volume: GLS model with restricted cubic splines. Model-predicted left and right ChP volume across each predictor in the spline model: age, estimated total intracranial volume, Talairach x-axis rotation, image resolution, scanner manufacturer, and sex. Upper panels, left hemisphere; lower panels, right hemisphere. Shaded bands and horizontal bars indicate 95% confidence intervals.

**Fig 2.**
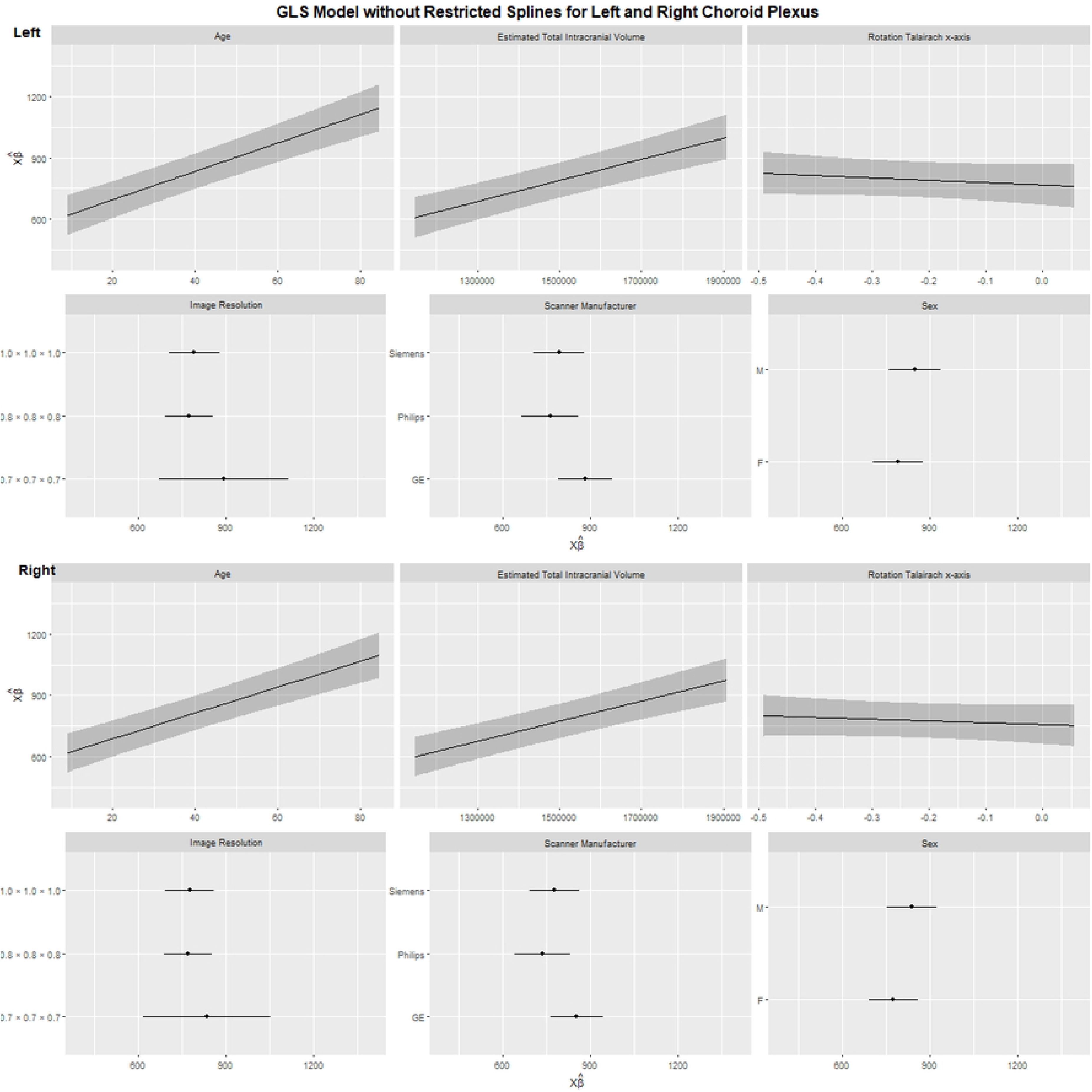
Choroid plexus volume: GLS model without restricted cubic splines. Model-predicted left and right ChP volume across each predictor in the linear model: age, estimated total intracranial volume, Talairach x-axis rotation, image resolution, scanner manufacturer, and sex. Upper panels, left hemisphere; lower panels, right hemisphere. Shaded bands and horizontal bars indicate 95% confidence intervals.

**Fig 3.**
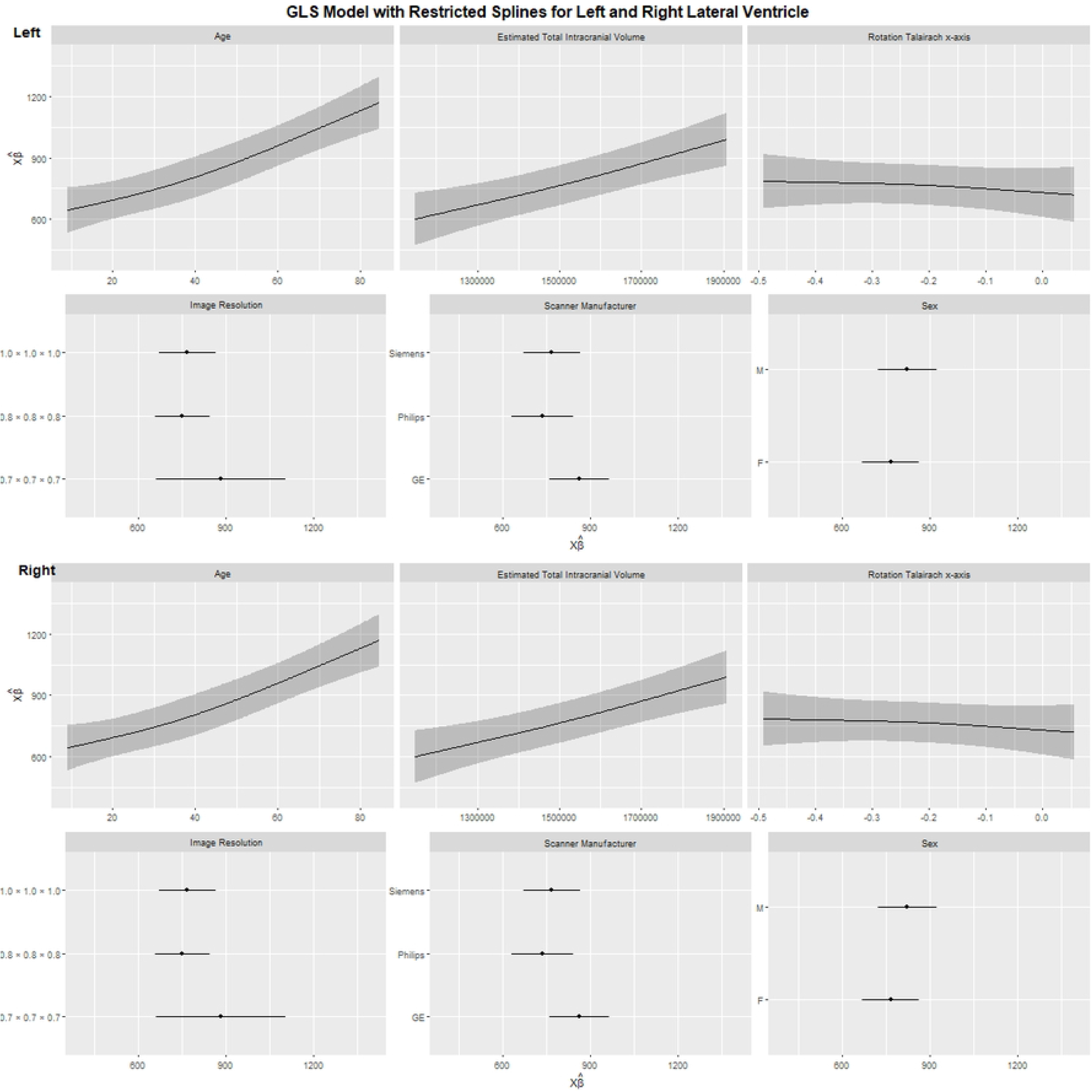
Lateral ventricle volume: GLS model with restricted cubic splines. Model-predicted left and right LV volume across each predictor in the spline model: age, estimated total intracranial volume, Talairach x-axis rotation, image resolution, scanner manufacturer, and sex. Upper panels, left hemisphere; lower panels, right hemisphere. Shaded bands and horizontal bars indicate 95% confidence intervals.

**Fig 4.**
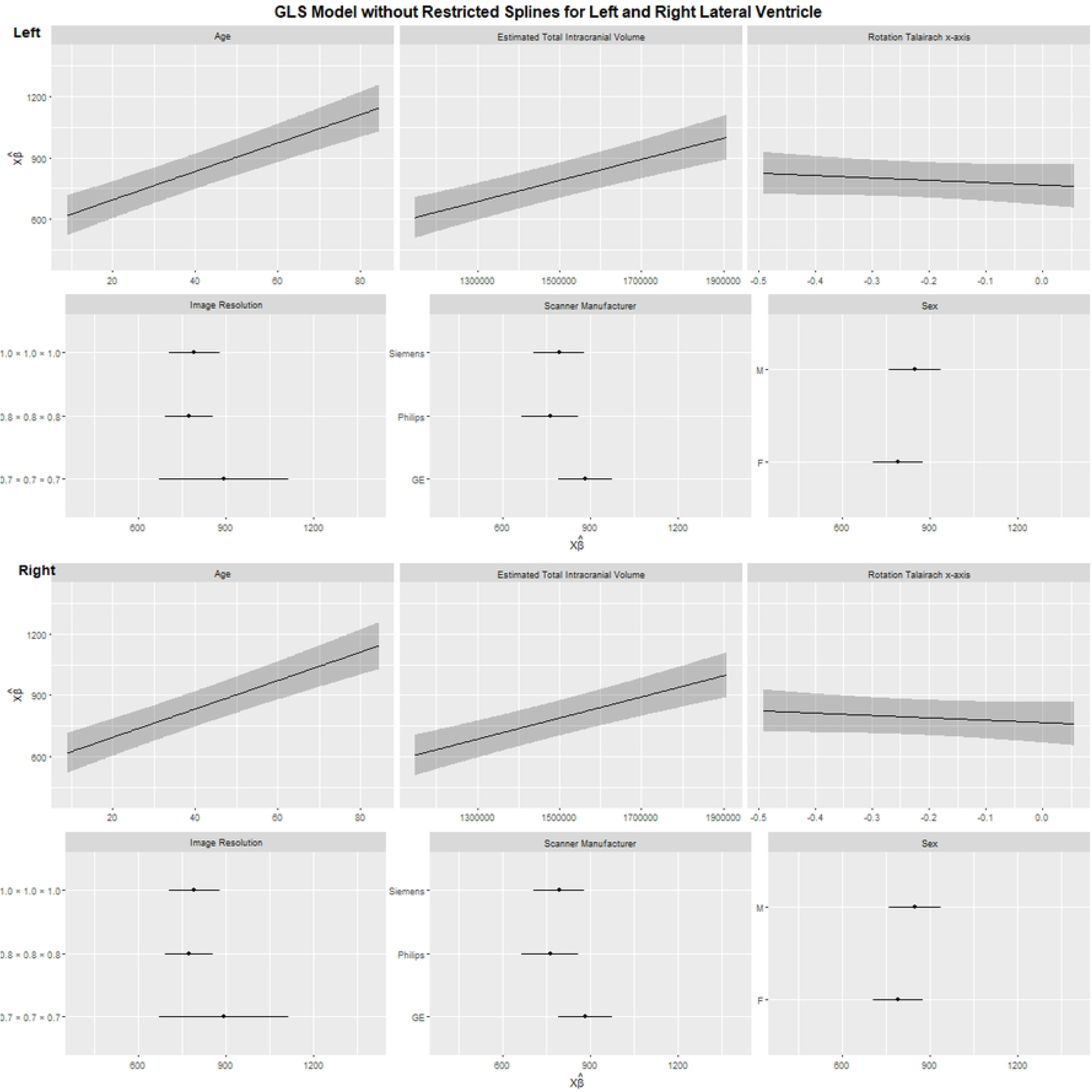
Lateral ventricle volume: GLS model without restricted cubic splines. Model-predicted left and right LV volume across each predictor in the linear model: age, estimated total intracranial volume, Talairach x-axis rotation, image resolution, scanner manufacturer, and sex. Upper panels, left hemisphere; lower panels, right hemisphere. Shaded bands and horizontal bars indicate 95% confidence intervals.

In the spline model, adjusting for all other covariates, males had 56.26 mm³ greater (95% CI: 4.35–108.2; p=0.034) average left ChP and 65.23 mm³ greater (95% CI: 15.99–114.5; p=0.009) average right ChP volume compared to females (Fig 1, Table 3). Sex differences in LV volume were not significant (left: p=0.825; right: p=0.500). These findings were also reflected in the linear model.

The three scanner manufacturers are significantly different in the left LV volume (p=0.019) in the spline model (Fig 3, Table 3). In a post hoc pair-wise comparison, the significant finding is between Philips and GE, where the average left LV is 2,024.32 mm³ higher in GE than Philips (95% CI: 592.88–3,455.76, p=0.006). Likewise, this is also the case between scans of the three image resolutions for the left LV volume (p=0.012). In a post hoc pair-wise comparison, the significant finding is between 0.8 mm and 1.0 mm, where average left LV volume is 1,535.76 mm³ higher in 1.0 mm scans than in 0.8 mm scans (95% CI: 433.12–2,638.40, p=0.006). In contrast, neither scanner manufacturer nor image resolution was associated with left or right ChP volume, or with right LV volume. Talairach x-axis rotation (rot_tal_x) was not associated with any outcome, whether modeled as a linear or nonlinear term.

Left and right ChP volumes were both significantly related to eTIV (p<0.001), with no significant nonlinear component (p=0.662 and p=0.638, respectively). The same pattern held for left and right LV volume (linear: both p<0.001; nonlinear: p=0.184 and p=0.180, respectively), in contrast with the age-LV relationship, there was no evidence of a nonlinear eTIV-LV relationship. Based on the linear model, a 100,000 mm³ difference in eTIV was associated with an average difference in left ChP volume of 51.63 mm³ (95% CI: 36.22–67.04), in right ChP volume of 49.22 mm³ (95% CI: 34.58–63.86), in left LV volume of 1,213.60 mm³ (95% CI: 978.26–1,448.90); and in right LV volume of 1,129.40 mm³ (95% CI: 923.69–1,335.0).

In the sex interaction model, the left and right ChP volumes were both related to age (p<0.001), this association was not significantly different between males and females (p=0.411, 0.060). LV volume was also related to age, and this association did significantly differ between males and females (p=0.002, 0.006). For males, the left LV increases by 2,138.80 mm³ (95% CI: 1845.55–2432.05) and the right LV increases by 1786.55 mm³ (95% CI: 1532.40–2040.70) per 10-year increase in age. For females, the left LV increases by 1679.10 mm³ (95% CI: 1400.40–1957.80) and the right LV increases by 1425.90 mm³ (95% CI: 1184.65–1667.25) per 10-year increase in age. At the sample median age and eTIV, males had greater right ChP volume than females by 57.44 mm³ (95% CI: 7.96 to 106.93), whereas the sex differences in left LV volume (females greater by 260.67 mm³; 95% CI: −532.16 to 1,053.50) and right LV volume (females greater by 354.27 mm³; 95% CI: −329.69 to 1,059.00) were not significant at that point. The overall tests of sex, which include the age×sex and sex×eTIV interaction terms, were significant for right ChP (p=0.009), left LV (p=0.017), and right LV (p=0.03) volume; for LV, this reflects the sex difference in the age slope described above rather than a constant offset. The overall test of sex was not significant for left ChP volume (p=0.088). There were no significant findings between scanner manufacturer, image resolution or rot_tal_x in any of the four regions. Left and right ChP volumes were both significantly related to eTIV (p<0.001), this association was not significantly different between males and females (p=0.273, 0.242). This was also the case for the left and right LV volume, which were related to eTIV (p<0.001), but were not significantly different between males and females (p=0.491, 0.387).

### Exploratory analyses of cognition

The estimated β coefficients, 95% confidence intervals, and p values for cognitive outcomes are reported in Table 6; all p values are uncorrected for multiple comparisons.

**Table 6.** Linear GLS models of cognitive outcomes on left and right ChP and LV volumes.

|  | Left ChP |  |  |  |
|---|---|---|---|---|
|  | β coef | Lower 95% CI | Upper 95% CI | p |
| PENN-40 | 0.0017 | 0.0003 | 0.0030 | **0.013** |
| FLANKER | -0.0045 | -0.0086 | -0.0004 | **0.032** |
| DCCS | -0.0005 | -0.0049 | 0.0039 | 0.821 |
| LSWMT | -0.0067 | -0.0127 | -0.0006 | **0.031** |
| PSM | -0.0078 | -0.0147 | -0.0009 | **0.026** |
| TPVT | -0.0030 | -0.0082 | 0.0022 | 0.253 |
|  | Right ChP |  |  |  |
|  | β coef | Lower 95% CI | Upper 95% CI | p |
| PENN-40 | 0.0013 | -0.0001 | 0.0027 | 0.065 |
| FLANKER | -0.0056 | -0.0099 | -0.0013 | **0.011** |
| DCCS | -0.0011 | -0.0057 | 0.0035 | 0.630 |
| LSWMT | -0.0039 | -0.0103 | 0.0026 | 0.239 |
| PSM | -0.0084 | -0.0157 | -0.0012 | **0.022** |
| TPVT | -0.0010 | -0.0065 | 0.0045 | 0.715 |
|  | Left LV |  |  |  |
|  | β coef | Lower 95% CI | Upper 95% CI | p |
| PENN-40 | 0.0001 | 0.0000 | 0.0002 | **0.026** |
| FLANKER | -0.0002 | -0.0005 | 0.0000 | 0.060 |
| DCCS | -0.0001 | -0.0003 | 0.0002 | 0.607 |
| LSWMT | 0.0001 | -0.0002 | 0.0005 | 0.544 |
| PSM | -0.0003 | -0.0007 | 0.0002 | 0.232 |
| TPVT | 0.0001 | -0.0002 | 0.0004 | 0.629 |
|  | Right LV |  |  |  |
|  | β coef | Lower 95% CI | Upper 95% CI | p |
| PENN-40 | 0.0001 | 0.0000 | 0.0002 | 0.249 |
| FLANKER | -0.0004 | -0.0007 | -0.0001 | **0.014** |
| DCCS | -0.0001 | -0.0005 | 0.0002 | 0.390 |
| LSWMT | 0.0001 | -0.0004 | 0.0005 | 0.735 |
| PSM | -0.0004 | -0.0009 | 0.0001 | 0.141 |
| TPVT | 0.0002 | -0.0002 | 0.0005 | 0.386 |

Note: ChP, choroid plexus; β coef, β coefficient; CI, confidence interval; LV, lateral ventricle; refer to Table 1 for full names of the cognitive measures. β coefficients are per 1 mm<sup>3</sup> of volume; models are adjusted for age, sex, scanner manufacturer, image resolution (omitted from the PENN-40 models), rot\_tal\_x, and eTIV. p values are not corrected for multiple comparisons. Bold indicates statistical significance.

### Penn Emotion Recognition Task (PENN-40)

Total PENN-40 score was significantly positively associated with left ChP volume (p=0.013) and left LV volume (p=0.026). On average, adjusting for all other covariates, a 100 mm³ difference in left ChP volume was associated with a 0.166 point difference in PENN-40 score (95% CI: 0.035–0.298); a 1,000 mm³ difference in left LV volume was associated with a 0.092 point difference in PENN-40 score (95% CI: 0.011–0.172).

### NIH Toolbox Related Cognition

Left ChP volume was significantly negatively associated with the FLANKER unadjusted scaled score (p=0.032), the LSWMT uncorrected standard score (p=0.031) and the PSM unadjusted scaled score (p=0.026). Each 100 mm³ difference in left ChP volume was associated with an average difference of −0.450 points in FLANKER (95% CI: −0.862 to −0.038), −0.668 points in LSWMT (95% CI: −1.274 to −0.063), and −0.780 points in PSM (95% CI: −1.467 to −0.093). Right ChP volume was significantly negatively associated with the FLANKER unadjusted scaled score (p=0.011) and the PSM unadjusted scaled score (p=0.022): average FLANKER score was 0.560 points lower (95% CI: −0.992 to −0.128) and PSM score was 0.845 points lower (95% CI: −1.570 to −0.120) comparing groups with a 100 mm³ difference in right ChP volume. Right LV volume was significantly negatively associated with FLANKER unadjusted scaled score (p=0.014), with an average 0.380 points lower (95% CI: −0.686 to −0.076) per 1000 mm³ difference. There were no other significant associations involving ChP or LV.

## Discussion

In this study, we manually segmented the lateral ventricle ChP in 700 MRI scans (699 analyzed after one processing failure) drawn from 35 site categories, spanning a broad range of age, sex, and scanner manufacturer; to our knowledge, this is the largest manually segmented ChP dataset assembled to date across this degree of demographic and technical heterogeneity. Consistent with our a priori hypotheses, both age and sex were significantly associated with ChP volume, with ChP volume tending to be larger in older individuals and showing a significant male-greater pattern, adjusting for all other covariates. There was no significant evidence of a non-linear relationship between age and ChP volume. LV volume, by contrast, showed a significant nonlinear relationship with age and no significant sex difference. Both ChP and LV volume scaled with eTIV, as expected. Scanner manufacturer and image resolution were significantly associated with left LV volume but not with ChP volume in either hemisphere, and the quality-control metric (Talairach registration rotation) was not significantly associated with any outcome. In exploratory analyses, larger ChP volume was nominally associated with worse inhibitory control/attention (FLANKER), episodic memory (PSM), and working memory (LSWMT; left only), but better emotion recognition (PENN-40) performance, whereas LV volume showed fewer associations (left LV with PENN-40; right LV with FLANKER). Together, these findings both replicate and extend prior, smaller-scale reports, and directly address the demographic and technical gaps identified in the introduction.

The positive relationship observed between age and ChP volume is consistent with a growing and large-scale literature, though the segmentation method underlying each study should be weighed alongside its findings. Two very recent large cohort studies converge on the same direction of effect: a UK Biobank analysis of 45,306 participants found ChP volume positively correlated with age (r=0.49, p<0.001) [30]; and a 2,137-participant Japanese cohort similarly found ChP volume increasing with age (r=0.351, p<0.001) [31]. Notably, the UK Biobank study segmented the ChP using FreeSurfer’s automated pipeline, and because FreeSurfer-derived ChP volumes have been shown to agree poorly with manual tracing across multi-vendor samples (Dice ≈0.29) [3], this association, and others like it that rely on FreeSurfer, may not fully reflect the true ChP-age relationship and should be interpreted with that caveat in mind. These findings echo smaller MRI cohorts (n=155, FreeSurfer with manual correction/QC [12]; HCP-Aging, n=641, Bayesian Gaussian mixture model [13]; n=98, deep learning validated against manual ground truth [11]), several of which also reported this relationship as nonlinear; the lack of evidence of nonlinearity in our analysis may reflect the substantially wider age range, which could smooth out curvature that is more detectable within a narrower adult-only window. We similarly found that LV volume related to age nonlinearly, consistent with well-established patterns of accelerating ventricular enlargement with advancing age, and that both ChP and LV volume scaled with eTIV, an expected relationship given that intracranial volume reflects overall brain and ventricular system size across development and aging. Taken together with a recent systematic review and meta-analysis linking ChP enlargement to Alzheimer’s disease [32], which, like most pooled analyses in this literature, aggregates studies using a mix of manual and automated segmentation methods, the results herein reinforce that age-related ChP growth is a robust, clinically relevant phenomenon that is best characterized with manually segmented, method-consistent data across a wide range of ages, which this dataset is uniquely positioned to provide. Because manual segmentation at this scale remains labor-intensive, our group is also developing a deep learning-based ChP segmentation tool trained on DataPlex’s manually segmented scans; by using manual tracing rather than FreeSurfer’s labels as ground truth, we anticipate this tool will achieve substantially higher accuracy than currently available automated methods, enabling these demographic relationships to be characterized in future cohorts far larger than manual segmentation alone can feasibly cover.

The current study found that males had greater ChP volume than females, with no significant sex difference in LV volume. This finding aligns with Eisma et al. (2024) [11], who reported significantly larger ChP volume in males across three MRI sequences, but contrasts with Alisch et al. (2021) [12], who found no significant cross-sectional sex difference, and with Zeng et al. (2024) [7], whose male-greater effect disappeared after covarying age and intracranial volume, a covariate we also included, which makes the persisting sex difference notable. The cross-sectional finding of greater male ChP volume is not necessarily inconsistent with Martinkova et al. (2023) [14], who reported that women’s ChP grows roughly twice as fast annually as men’s in a longitudinal ADNI sample; a steeper female growth trajectory and a male-larger cross-sectional average can coexist at different points across the lifespan. A plausible biological substrate for sex-specific ChP characteristics has recently emerged at the molecular level: a 2025 cross-species study found pronounced, conserved sex differences in ChP secretory and immune signaling, a female-biased immune and antigen-presentation profile versus a male-biased complement/coagulation and ion-transport profile, replicated in both mouse and human ChP transcriptomic and CSF proteomic data [33]. While this work characterizes functional and cellular differences rather than gross volume directly, it offers a credible mechanistic basis for sex-divergent ChP biology that merits further investigation.

Scanner manufacturer and image resolution were significantly associated with left LV volume but not with ChP volume (in either hemisphere) or right LV volume, and Talairach rotation was unrelated to any outcome. The ChP scanner findings were, however, model-dependent: scanner associations with ChP volume were borderline in the primary model (p=0.069 and p=0.095), and in the sensitivity model substituting GM signal-to-noise ratio for Talairach rotation (S1 File), scanner manufacturer was significantly associated with left and right ChP volume (larger in GE than in Philips and Siemens scans), and higher GM signal-to-noise ratio was associated with smaller right ChP volume. This pattern is notable against the backdrop of the broader volumetric MRI reliability literature, which has long established that vendor and acquisition parameters systematically bias structural and ventricular measurements [18,19], and against more recent work showing that automated (FreeSurfer-based) ChP segmentation specifically is highly scanner-sensitive, agreeing poorly with manual ground truth across multi-vendor samples (Dice ≈0.29 [3]) and only moderately even within a single scanner (r=0.659 [16]). The relative, though incomplete, scanner-robustness of the ChP measurements, despite being drawn from 35 heterogeneous site categories, may reflect the value of manual segmentation as the reference-standard method: where automated pipelines absorb scanner-related noise directly into the measurement, trained raters tracing ChP boundaries by eye may be comparatively insulated from potential scanner or QC metric related biases. That LV volume, derived in part from automated FastSurfer output in the corrected-volume calculation, retained scanner sensitivity is consistent with this interpretation and with the established automated-volumetry reliability literature. Combined with the broader push toward explicit site/scanner modeling in pooled multi-site neuroimaging work, the results herein support treating manual ChP segmentation, paired with explicit scanner-level covariates, as a methodologically sound approach for pooling heterogeneous public datasets.

In exploratory analyses that were not corrected for multiple comparisons, ChP and LV volumes showed nominal associations with several cognitive domains, and the pattern was not a uniform “larger volume, worse cognitive performance” relationship. Greater left ChP and left LV volume were nominally associated with better PENN-40 emotion recognition performance, whereas greater left and right ChP volume was nominally associated with worse inhibitory control/attention (FLANKER) and worse episodic memory (PSM) and working memory (LSWMT, left ChP only), and greater right LV volume was associated with worse inhibitory control/attention. The negative associations are directionally consistent with the existing ChP-cognition literature, where ChP enlargement has repeatedly been linked to worse cognitive outcomes, in psychosis, where larger ChP volume correlates with worse composite cognition and elevated interleukin-6 [1]; in schizophrenia-spectrum disorders, where ChP-defined ventricular subgroups show distinct cognitive profiles [16]; and, most recently, at population scale, where greater ChP volume predicted faster cognitive decline and higher incident dementia risk in the UK Biobank [30]. The positive PENN-40 association runs counter to this pattern and is harder to reconcile with the existing literature; we are aware of no comparable study linking larger ChP volume to better performance on any cognitive measure. Because both ChP volume and emotion recognition performance increase from childhood into adulthood, residual age-related confounding across this wide age range is a plausible explanation. Given the number of tests, unequal battery coverage across studies, and the wide age range, these associations should be regarded as hypothesis-generating. Replication in a sample with more uniform cognitive battery coverage, and ideally with a targeted hypothesis about emotion recognition specifically, would help clarify whether the PENN-40 finding reflects a genuine domain-specific or developmental-stage effect, or a feature of this subsample.

Several additional limitations should be considered. First, this study is cross-sectional, precluding causal or within-person longitudinal inference about how ChP and LV volume change with age. Second, despite DataPlex’s breadth relative to prior ChP cohorts, the sample’s age range (6.3 to 89.8) does not extend into infancy or early childhood, a developmental window in which the ChP undergoes distinct, rapid structural change tied to the establishment of CSF production [5], a window that existing pediatric ChP literature has so far treated as a separate line of inquiry from adult and aging-focused work. Third, DataPlex pools scans from 35 site categories with heterogeneous original recruitment criteria, including both healthy and clinical populations (e.g., early psychosis, clinical-high-risk, and epilepsy cohorts); phenotype was a selection criterion for dataset diversity but was not modeled as a covariate in the present analyses, and residual clinical heterogeneity could confound demographic and cognitive associations alike. In addition, the 0.7 mm resolution group was small (n=22) and came from a single site (Human Connectome Project Young Adult, Siemens), so estimates for this resolution are confounded with age group, study, and scanner manufacturer. Fourth, cognitive battery availability varied substantially by contributing study and also within those studies, by differences in exam acquisition, limiting statistical power and generalizability for the cognitive findings discussed above. Fifth, corrected LV volume estimate depends on an automated FastSurfer-derived calculation, which may propagate residual segmentation error despite correction for ChP mislabeling. Sixth, although manual segmentation is the reference-standard method, it still depends on rater judgment; despite a mean inter-rater Dice coefficient of 0.81, some residual measurement variability across 10 raters is possible, and finer-grained technical heterogeneity (scanner model, software version, head coil channel count) could not be modeled due to inconsistent availability across contributing studies.

## Conclusion

In this study, we analyzed DataPlex, a manually segmented lateral ventricle choroid plexus dataset of 700 MRI scans spanning a wide range of age, sex, and scanner types and, to our knowledge, the largest and most demographically diverse manually segmented ChP resource assembled to date. We found that age and sex were significantly associated with ChP volume, that ChP volume was comparatively robust to scanner manufacturer and image resolution relative to lateral ventricle volume, and that ChP and lateral ventricle volume showed exploratory associations with several cognitive domains. These findings underscore the importance of accounting for demographic and technical covariates in future choroid plexus research and establish reference benchmarks that automated and semi-automated segmentation methods, including the deep learning tool our group is developing from this dataset, can be validated against going forward.

## Data availability

The manual segmentations are uploaded on the NDA (Collection ID# 5175). This data containing manual segmentations will be released by the NDA within 4 weeks. The corresponding T1s are from a mix of open source and data use agreement scans (see S1 File for data acquisition details), details on which scan from the project was used can be derived from the NDA’s image03.txt file, in the *src_subject_id* column. Questions regarding data availability can be sent by email to the corresponding author.

## Acknowledgments

We thank the trained raters who performed the manual choroid plexus segmentations (D. Ying, D. P. Dhunna, E. Oss, M. Hricay, V. Lyons, A. Adesokan, W. Molho, B. Stiltner, P. Patel, and N. Tolentino); L.-Y. Huang, N. Raymond, and V. Zeng served as expert quality-control raters. We also thank Oshin Kushwaha for assistance in setting up the project infrastructure.

This study used data from multiple projects, and we thank the principal investigators of these projects for making their data available and the participants that were enrolled in those projects. The principal investigators of these projects did not participate in the analysis or writing of this manuscript. Acknowledgments required by the data use agreements of the contributing studies follow.

## ABCD

Data used in the preparation of this article were obtained from the Adolescent Brain Cognitive Development (ABCD) Study (https://www.nbdc-datahub.org/), held in the NIMH Data Archive (NDA). This is a multisite, longitudinal study designed to recruit more than 10,000 children age 9–10 and follow them over 10 years into early adulthood. The ABCD Study® is supported by the National Institutes of Health and additional federal partners under award numbers U01DA041048, U01DA050989, U01DA051016, U01DA041022, U01DA051018, U01DA051037, U01DA050987, U01DA041174, U01DA041106, U01DA041117, U01DA041028, U01DA041134, U01DA050988, U01DA051039, U01DA041156, U01DA041025, U01DA041120, U01DA051038, U01DA041148, U01DA041093, U01DA041089, U24DA041123, and U24DA041147. A full list of supporters is available at https://abcdstudy.org/federal-partners.html. A listing of participating sites and a complete listing of the study investigators can be found at https://abcdstudy.org/consortium_members/. ABCD consortium investigators designed and implemented the study and/or provided data but did not necessarily participate in the analysis or writing of this report. This manuscript reflects the views of the authors and may not reflect the opinions or views of the NIH or ABCD consortium investigators. The ABCD data repository grows and changes over time. The ABCD data used in this report came from doi:10.15154/z563-zd24.

## AMP SCZ

Data used in the preparation of this article were obtained from the AMP SCZ study (https://www.ampscz.org), held in the NIMH Data Archive (NDA). This is a multisite, longitudinal study designed to recruit more than 2,500 subjects ages 12 to 30 and follow them over the course of 24 months. The AMP SCZ research program is supported by contributions from the AMP SCZ public and private partners, including the National Institute of Mental Health (NIMH) under award numbers U24MH124629, U01MH124631, and U01MH124639, the Wellcome Trust (220664/Z/20/Z and 220664/A/20/Z), and the NHMRC CRE PRE-EMPT (1198304). A full list of supporters is available at https://www.ampscz.org/about/partners/, and a complete listing of the study investigators is at https://www.ampscz.org/about/research-networks/. AMP SCZ consortium investigators designed and implemented the study and/or provided data but did not necessarily participate in the analysis or writing of this report. This manuscript reflects the views of the authors and may not reflect the opinions or views of the NIH or AMP SCZ consortium investigators.

## HCP Young Adult

Data were provided in part by the Human Connectome Project, WU-Minn Consortium (Principal Investigators: David Van Essen and Kamil Ugurbil; 1U54MH091657) funded by the 16 NIH Institutes and Centers that support the NIH Blueprint for Neuroscience Research; and by the McDonnell Center for Systems Neuroscience at Washington University.

## HCP Aging

Research reported in this publication was supported by the National Institute on Aging of the National Institutes of Health under Award Number U01AG052564. The content is solely the responsibility of the authors and does not necessarily represent the official views of the National Institutes of Health. The HCP-Aging 2.0 Release data used in this report came from doi:10.15154/1520707.

## HCP Development

Research reported in this publication was supported by the National Institute of Mental Health of the National Institutes of Health under Award Number U01MH109589 and by funds provided by the McDonnell Center for Systems Neuroscience at Washington University in St. Louis. The HCP-Development 2.0 Release data used in this report came from doi:10.15154/1520708.

## SRPBS

Data used in the preparation of this work were obtained from the DecNef Project Brain Data Repository (https://bicr-resource.atr.jp/srpbsopen/), collected as part of the Japanese Strategic Research Program for the Promotion of Brain Science (SRPBS) supported by the Japanese Advanced Research and Development Programs for Medical Innovation (AMED).

## Other contributing studies

Oxytocin: UL1-TR000064, R01AG059809, R24AG039350, R24AG054355, 3R01AG072658-01A1S, and T32AG020499; National Science Foundation Cooperative Agreement No. DMR-1157490; U.S. Navy Office of Naval Research Grant N00014-21-1-2201. NAPLS3 (doi:10.15154/304r-9v58): U01MH081928, U01MH081988, U01MH081984, U01MH082004, U01MH081902, U01MH082022, U01MH081944, U01MH076989, U01MH081857. BANDA (doi:10.15154/6fng-rj25): U01MH108168. OCD (doi:10.15154/aps3-0723): R01MH059299-15. Paranoia (doi:10.15154/pmb4-r907): R21MH112930. SPARCS: K01MH114012, R01MH107579-04. BICEPS: R01MH092440. EMBARC (doi:10.15154/e99n-np66): U01MH092221, U01MH092250. DLBS: R37AG-006265. DCAM (doi:10.15154/6neq-wm52): U01MH109991. PDC (doi:10.15154/00b0-hw43): U01MH110008. LateASD (doi:10.15154/nr0y-k674): R01MH103494. HCP-Epilepsy: U01NS093650. Diabetes-R01: R01AG070077. 3671: K23MH108711. 3854: R01MH126181. 3925: R21MH125044. SenDep (doi:10.15154/z3e6-w042): R01MH115953. OPTIMUM: R01MH114966, R01MH114969, R01MH114970, R01MH114980, R01MH114981. ABD: Natural Sciences and Engineering Research Council Discovery Grants (Canada). Late Life Depression: The First Affiliated Hospital of Nanchang University Internal Funding.

## Supporting information

**S1 File.** Supplementary information. Contributing studies and sample sizes (Table A), data use agreements, cognitive sample details, and the sensitivity analysis substituting gray matter signal-to-noise ratio for Talairach x-axis rotation (Table B, Figs A–D).

**S1 Checklist.** STROBE checklist for cross-sectional studies.

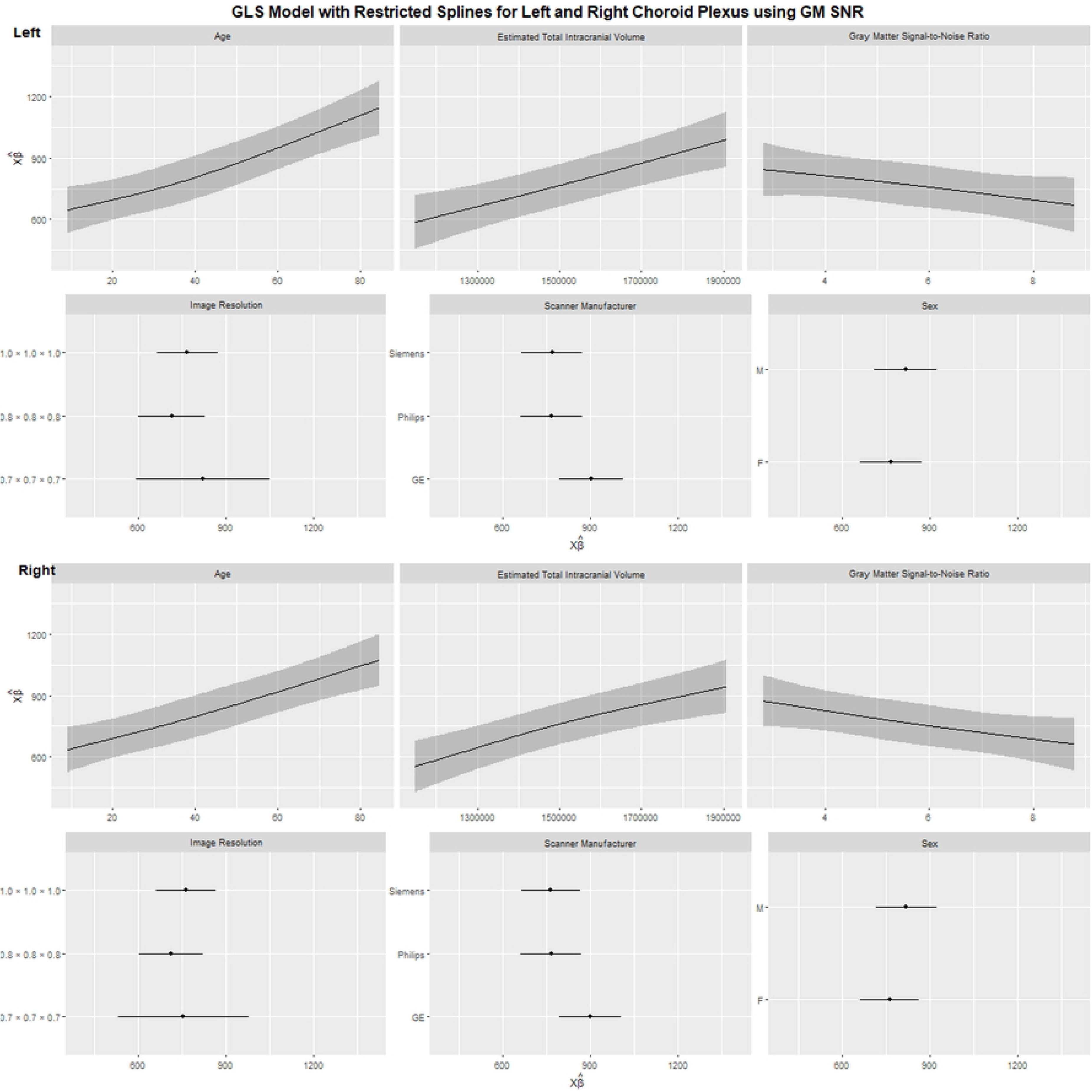

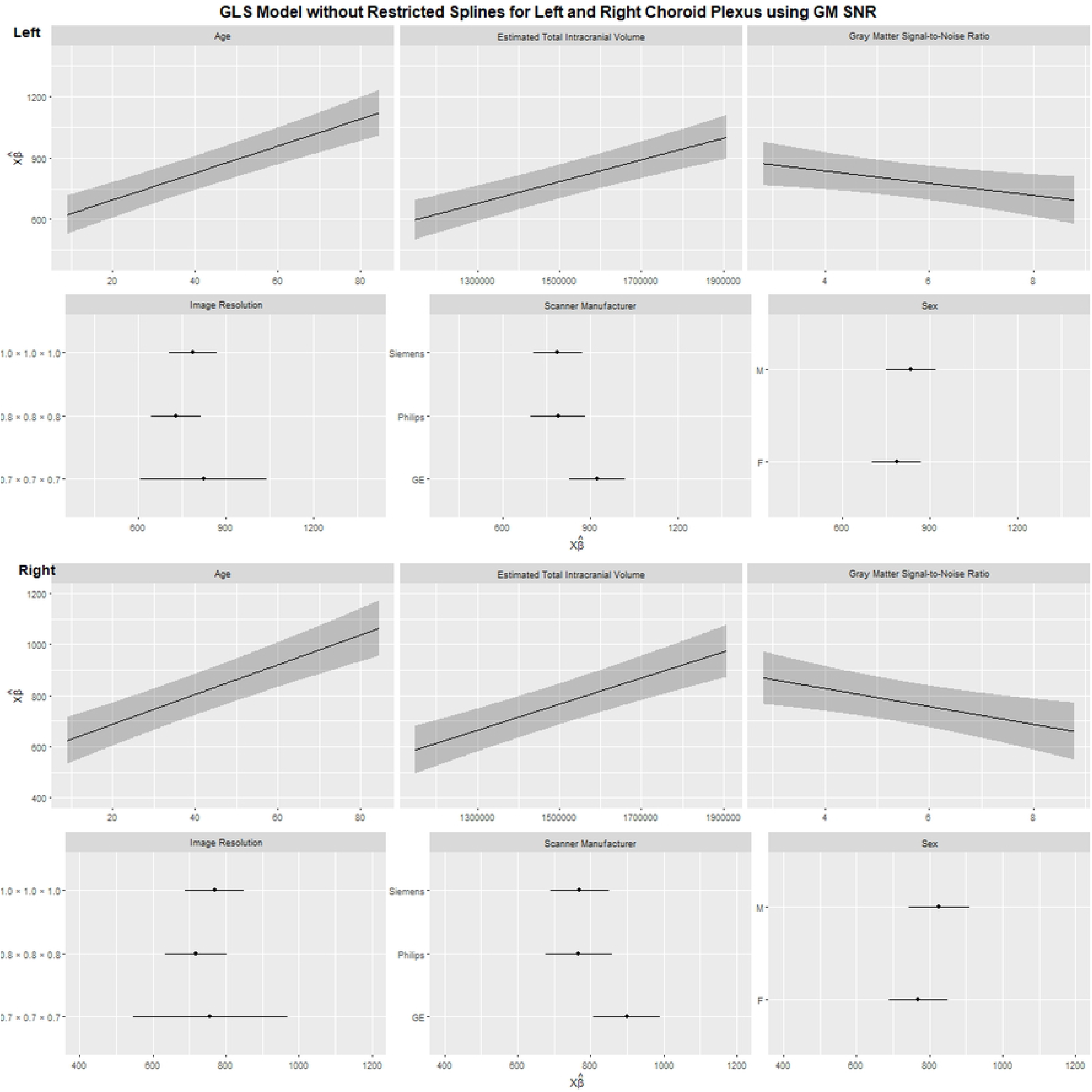

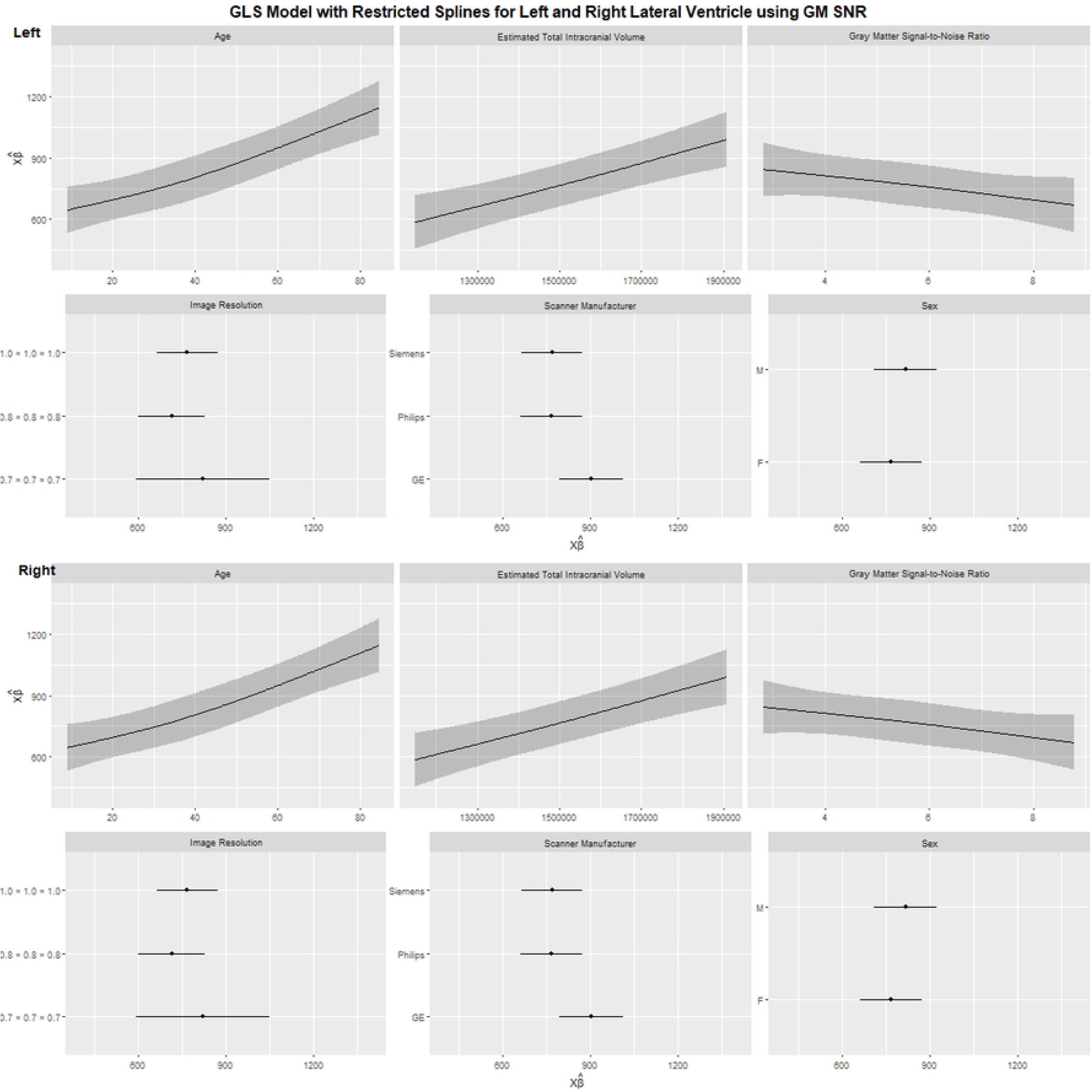

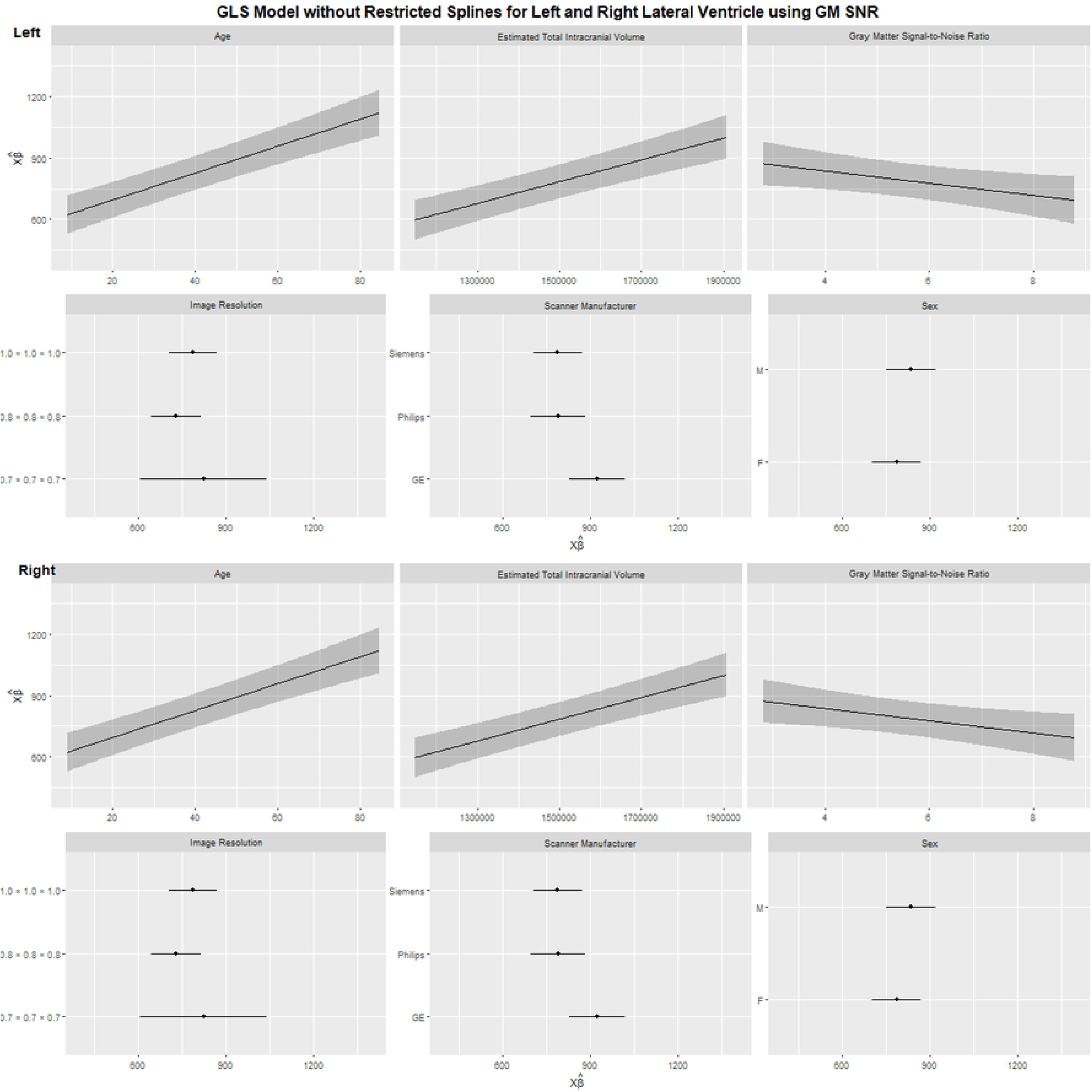

## Notes

### Competing Interest Statement

The authors have declared no competing interest.

